# From short-lived to persistent: The significance of extracellular RNA in disinfected drinking water microbiomes

**DOI:** 10.1101/2025.10.18.683149

**Authors:** Sakcham Bairoliya, Sudarsan Mugunthan, Dan Cheng, Yi-Nan Liu, Staffan Kjelleberg, Bin Cao

**Author notes:** Corresponding author: Dr. Bin Cao, Gene and Linda Voiland School of Chemical Engineering and Bioengineering, Washington State University, Pullman, WA Wegner Hall 309 (Office) and 325 (Lab).

## Abstract

To assure the safe distribution of drinking water, it is critical to identify adaptations that allow microorganisms to thrive in an oligotrophic system under disinfectant stress. Microbial adaptive responses are determined using RNA–based differential transcriptomics which cannot be performed in environmental settings due to lack of a baseline. This study introduces the concept of using extracellular RNA (eRNA) as a reference to determine important adaptive features in disinfected drinking water microbiome. Using this method, we determined energy efficiency, dynamic membrane composition, oxidative stress response, and proper protein folding to be extremely important for survival in drinking water systems. Additionally, multiple antibiotic resistance genes are expressed and upregulated in intracellular RNA fraction, indicating selection pressure. Non-coding RNA and transfer-messenger RNA found in eRNA suggest important regulatory functions. This study advances our understanding of eRNA in drinking water microbiomes and provides a framework for exploration in other environmental systems.

## Main text

The drinking water microbiome is extremely important to enable safe distribution of drinking water^1^. Molecular methods including DNA–based sequencing has been extensively used to illustrate the composition and functional potential of the drinking water microbiome^2–4^. However, most of these studies do not account for extracellular DNA, which can be present in significant proportions, especially in disinfected drinking water systems^5^.

RNA-based community detection has also been utilized to study the drinking water microbiome^6,7^. However, the drinking water metatranscriptome has not yet been characterized due to methodological challenges, including low biomass, low RNA extraction efficiency, a lack of comparative samples, and limited access to DWDS infrastructure^8^. It is essential to understand how these microorganisms thrive in a nutrient limited environment under disinfection stress. However, in-situ environmental RNA studies can only facilitate functional characterization of the microbiome due to the lack of a baseline for comparison^9^. Intriguingly, previous experimental findings using model organisms demonstrate that RNA from lysed cells can persist extracellularly for extended periods, offering valuable insights into cellular responses under disinfectant stress^10^. Extracellular RNA (eRNA) in drinking water may serve as a reference point for investigating the cellular mechanisms that support organism survival and proliferation.

We hypothesize that eRNA persists in the drinking water distribution system (DWDS) and that characterizing it will reveal crucial cellular decisions for survival at the transcriptional level. To investigate this, we integrated metagenomic and metatranscriptomic approaches and compare both extracellular nucleic acid (eNAC) and intracellular nucleic acids (iNAC) to provide a comprehensive understanding of DWDS microbiome. This comparison highlights the active community composition, preferred metabolic pathways, and expression of antibiotic resistance genes in monochloraminated DWDS.

### Comprehensive sampling of the drinking water microbiome

Drinking water was sampled directly from the tap using glass fiber membranes placed in inline filter holders connected to a 4-way flow splitter. Two consecutive samplings were conducted and a total of ∼40 L of drinking water per replicate was collected from the tap over a period of four hours (Figure 1A). Water samples were collected at the start (time 0), middle (time 2), and end (time 4) points of the sampling process and analyzed using a flow cytometer after live-dead staining^11,12^. The intact cell count decreased from 3 × 10^4^ (± 1.5 × 10^4^) cells/mL at time 0 to 3.2 × 10^3^ (± 0.9 × 10^3^) cells/mL at time 4 (Figure S1A, Supplementary Information (SI)). However, microbial phenotypic diversity (Figure S1B) increased throughout the sampling process. The phenotypic diversity at the beginning of sampling differed significantly from that at the end (Figure 1B), highlighting the dynamic nature of the drinking water microbiome and reflecting the thoroughness of the sampling process. Drinking water chemistry was only determined at the end of the sampling process, reflecting drinking water chemistry of actively disinfected water (Table S1).

**Figure 1.**
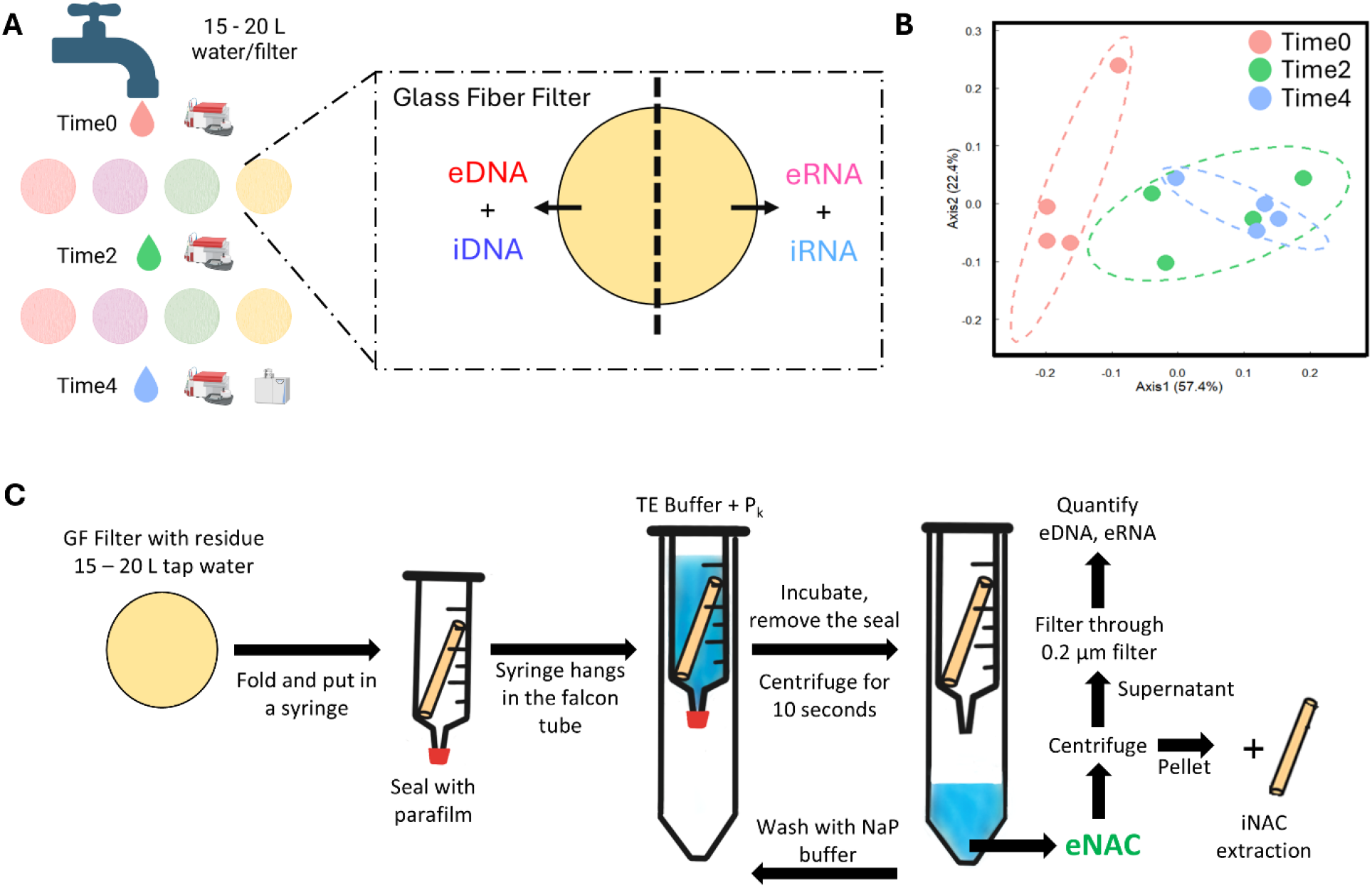
Study design. **A)** Schematic illustration of the sampling process to obtain different fractions of nucleic acids (eNAC + iNAC). **B)** Phenotypic diversity of drinking water before (time0), in-between (time2) and after (time4) the sampling process, **C)** Schematic illustration of the eNAC extraction process.

The four fractions of the intracellular and extracellular nucleic acids (iDNA, eDNA, iRNA, and eRNA) were extracted (Figure 1C, S2; Text S1), purified (Table S2), and sequenced as illustrated in Figure 1C. We could recover 56 ± 5 ng/L of eDNA which is close to our previous estimate from the same location^5^ while ∼140 ng/L of eRNA was recovered (Figure 2A).

**Figure 2.**
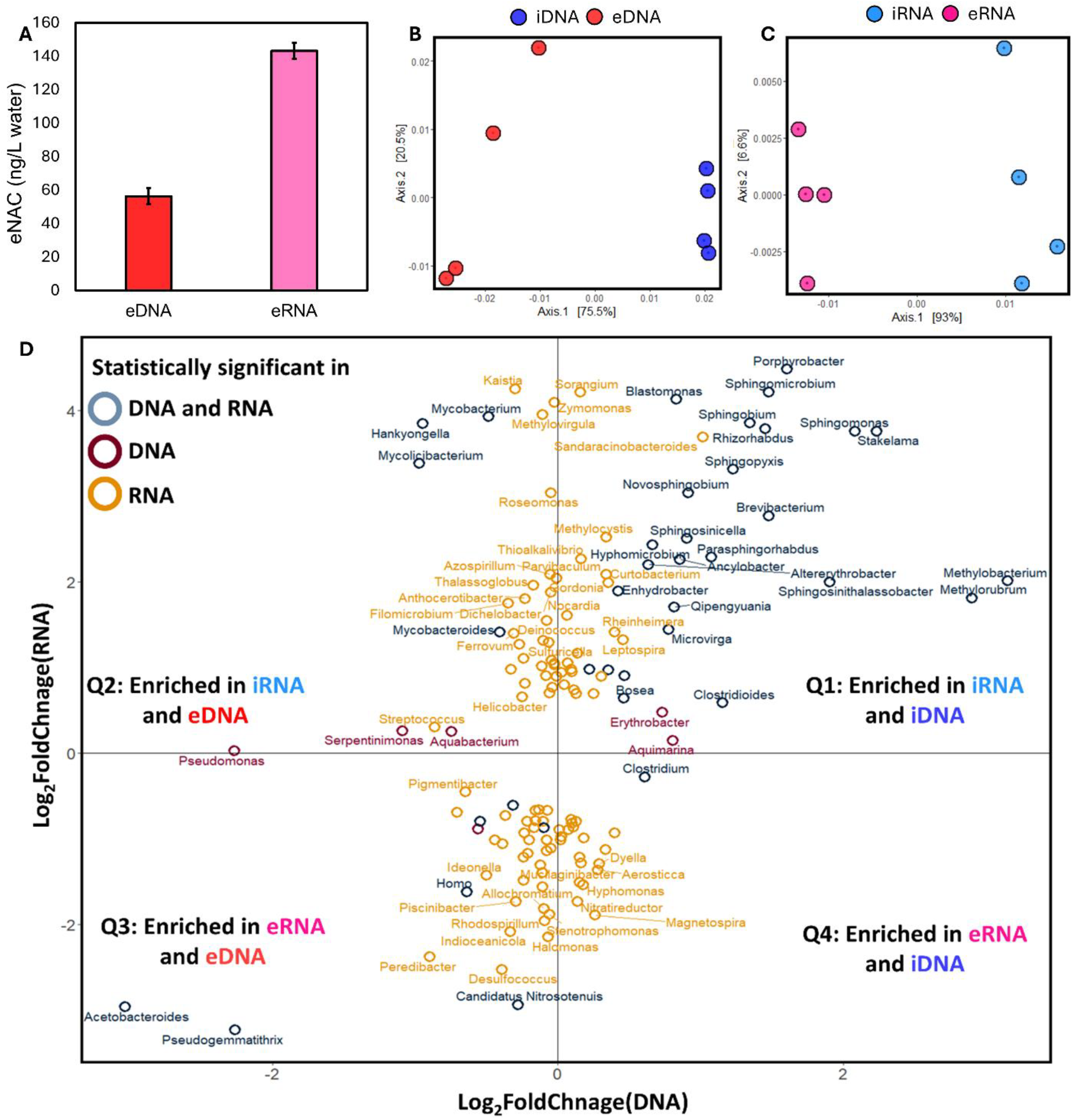
eNAC and its influence on detection of the drinking water community. **A)** Amount of extracellular DNA and RNA (eDNA and eRNA, respectively) extracted per liter of drinking water sampled (n = 4, error bar represents standard deviation). **B)** PCoA plot showing the difference between communities detected in eDNA and **C)** eRNA based on Bray-Curtis dissimilarity. **D)** Dynamics of drinking water community revealed via sequencing of iNAC and eNAC. All genera significantly different in RNA (basemean > 500, p.adjusted < 0.01) or DNA (basemean >50, p.adjusted < 0.01) were retained. Q1 represents genera which are live, metabolically active, and resistant to MCA. Q2 represents genera which are metabolically active, major part of biofilms in DWDS, and might be involved in active DNA release. Q3 represents taxa which are not resistant to MCA or does not originate from drinking water while Q4 contains taxa which are involved in active eRNA release. Different colors are used to indicate differentially abundant taxa in DNA and RNA, only DNA and only RNA.

### Distinct microbial communities are detected in each nucleic acid fraction

Approximately 55% of the total recovered sequences for DNA and around 99% for RNA were classified by Kraken2 with majority reads classified as bacteria (Table S3). eDNA and iDNA exhibited a comparable number of detected genera (observed richness, (Figure S3A)). Previous studies have shown a decrease in α-diversity indices following the removal of eDNA^5,7^, suggesting that eDNA may lead to false positive detection of genera. Similar patterns were observed with iRNA, which displayed a significantly greater number of detected genera compared to eRNA (p < 0.001, Figure S3B).

The community identified based on the extracellular fraction differed from that using the intracellular fraction (Figure 2B, C). The results were corroborated using a hierarchical clustering approach (Figure S3C, D). PERMANOVA results suggest that 73% and 91% of the variation in data can be explained by the fractions of nucleic acid (intracellular or extracellular) for DNA and RNA respectively (Table S4). The dispersion analysis reveals that the iDNA community was more consistent compared to the eDNA community while the opposite was observed for RNA (Figure S3E, F), confirming the contribution of eNAC to the variation in detected community composition. iDNA samples are distinct from eDNA samples because of differences in the abundance of *Methyloruburum, Methylobacterium, Sphingomonas* and *Pseudomonas* (Figure S4A). Similarly, *Sphingomonas, Methylobacterium, Nitrosomonas*, and *Saccharobesus* contribute to the differentiation between iRNA and eRNA samples (Figure S4B). A comparison of all four nucleic acid fractions reveals distinct microbial communities in each fraction (Figure S4C).

### Community composition from different NAC fractions reveals the activity of different genera

The 30 most abundant genera in each NAC fraction are detailed in the SI (Figures S5, S6). The fold change of taxa between eNAC and iNAC revealed significant differences in the detection of different genera (Figure 2D). Genera such as *Methylobacterium, Methylorubrum, Sphingomonas, Hyphomicrobium, Novosphingobium, Sphingobium, Rhizorhabdus*, and *Sphingopyxis* were enriched in iDNA and iRNA (Q1). These genera are commonly found in the DWDS, exhibit high resistance to MCA disinfection^2,13^, and thrive in the system. Quadrant 2 (Q2) includes genera enriched in the eDNA and iRNA fractions. Intriguingly, this group contains some hygienically relevant organisms such as *Mycobacterium, Mycolicibacterium, Mycobacteroides, Streptococcus*, and *Pseudomonas. Mycobacterium and Pseudomonas* are widely enriched in MCA treated distribution systems and are resistant to it^14^. Q2 taxa may also actively release eDNA, as part of biofilm formation.

One of the objectives of analyzing eNAC is to assess its role in detecting taxa that may not originate from the sampled ecosystem. For instance, nucleic acids from “homo”, indicating human contamination, are enriched in the extracellular fraction. This highlights the effectiveness of our method in distinguishing nucleic acid fractions. In Quadrant 3 (Q3), genera such as *Acetobacteroides. Pseudogemmatithrix, Candidatus Nitrosotenuis*, and *Halomonas*, were enriched in eDNA and eRNA, suggesting that these genera either did not thrive in the DWDS or their NAC did not originate from within the DWDS. If we were to analyze total NAC, as is commonly done in most studies, this distinction would not be possible. Quadrant 4 (Q4) features taxa such as *Clostridium* which is enriched in iDNA and eRNA. Q4 taxa may actively secrete eRNA.

### Metagenome shows high abundance of transposases, diverse ARGs and virulence factors

The general metagenome description can be found in Text S2. There was no substantial difference in the overall functions retrieved from the metagenome between iDNA and eDNA (Figure 3A). Among the top 30 COGs and KEGG Orthologs (KOs), 7 were classified as transposases (Table S5, S6). A total of 13868 different COGs/ EggNOG were identified in the metagenome, with 6,927 (having more than 20 counts across all samples) used for differential abundance analysis. We found 2,446 COGs that were differentially abundant between eDNA and iDNA (p-adjusted < 0.01). Among them, 1043 and 845 COGs were significantly enriched in iDNA and eDNA, respectively (p-adjusted < 0.01 and |log_2_(fold-change)| > 0.58, Figure 2B). Although eDNA did not impact the qualitative analysis of the DWDS metagenome, it obscured the quantitative assessment of various genes across different pathways, as evidenced by the differential abundance analysis.

**Figure 3.**
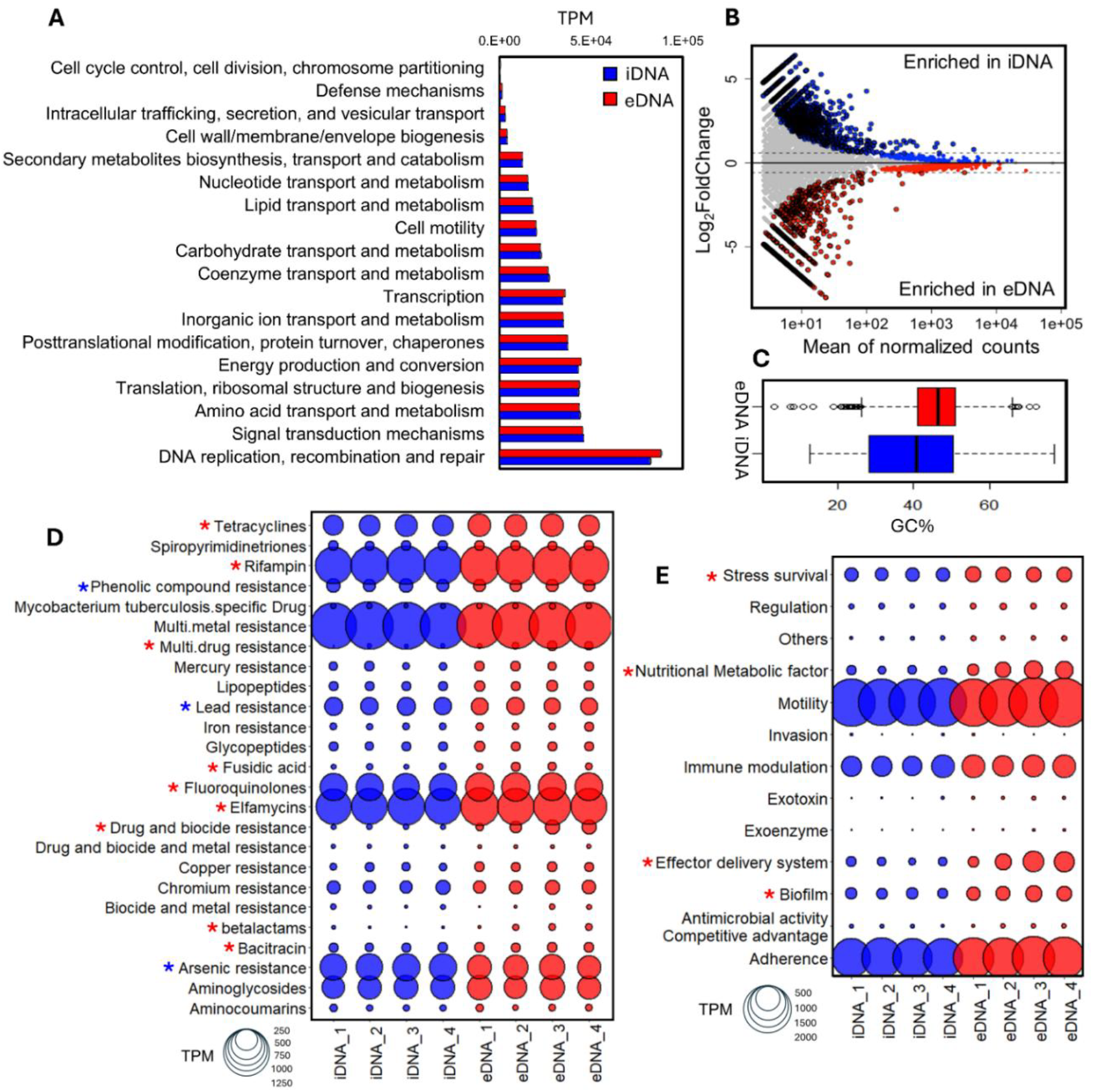
Features of the drinking water metagenome. **A)** General functions represented in the drinking water metagenomes by iDNA and eDNA according to COG categories. **B)** Fold change of different COGs. Significantly different COGs defined as COGs with p-adjusted < 0.01 and |log_2_(fold-change)| > 0.58. **C)** Mean GC content of coding regions enriched in iDNA and eDNA. **D)** Antibiotic, biocide and metal resistance genes and **E)** Virulence gene annotated in the drinking water metagenome using MEGARes^15^ and Virulence Factor database^16^. Statistically significant difference for **D)** and **E)** is indicated by ‘*’ (p < 0.05). ‘*’ indicates significant enrichment in iDNA and ‘*’ represents significant enrichment in eDNA. TPM: Transcripts per million.

In our previous work^10^, we found that persistence of eDNA is influenced by the GC content of the genes. This was verified using the current dataset. The coding regions enriched in eDNA exhibited an average GC content of 46%, while those enriched in iDNA had a mean GC content of 40% (Figure 3C), suggesting that genes with higher GC content are more likely to persist in the extracellular environment.

The metagenome harbors various antimicrobial resistance genes (ARGs) that confer resistance to a wide range of antibiotics, biocides, and heavy metals (Figure 3D). ARGs were found to be differentially abundant in eDNA and iDNA. Resistance genes for phenolic compounds and heavy metals such as lead and arsenic were found to be enriched in the iDNA, while genes associated with multi-drug resistance, rifampin-, betalactam-, tetracycline-, elfamycin-, and biocide-resistance were enriched in eDNA. Additionally, multiple virulence genes related to biofilm formation (alginate production, quorum sensing, and efflux pumps), stress tolerance (including detoxification of hydrogen peroxide and proteases), effector delivery system (secretion systems and iron acquisition), and nutritional metabolic factors were significantly enriched in eDNA (Figure 3E).

### Metatranscriptome shows different RNA subtypes in eRNA and expression of multiple antibiotic resistance genes

To reveal the actual gene expression patterns in the DWDS, we analyzed the metatranscriptome by aligning the RNA reads to the assembled metagenome and estimating the expression of coding regions (mapping rate = 99.7%). The types of RNA and small RNA (sRNA) were examined using Infernal^17^ and the Rfam database^18^ (Figure 4A). Majority of the RNA reads were assigned to the small and large subunits of rRNA for bacteria, eukarya, and archaea. Notably, eRNA contained a major portion of rRNA for eukarya and archaea. Additionally, tmRNA SsrA was enriched in the eRNA fraction^19^, along with RNase P, which is responsible for its processing. Several other small RNAs, such as 6S (which regulates sigma factor 70), poplar-1, ykkC-yxkD (guanidine-I riboswitch regulating efflux pumps and detoxification systems), and traJ-II (which regulates conjugation), were found to be enriched in iRNA.

**Figure 4.**
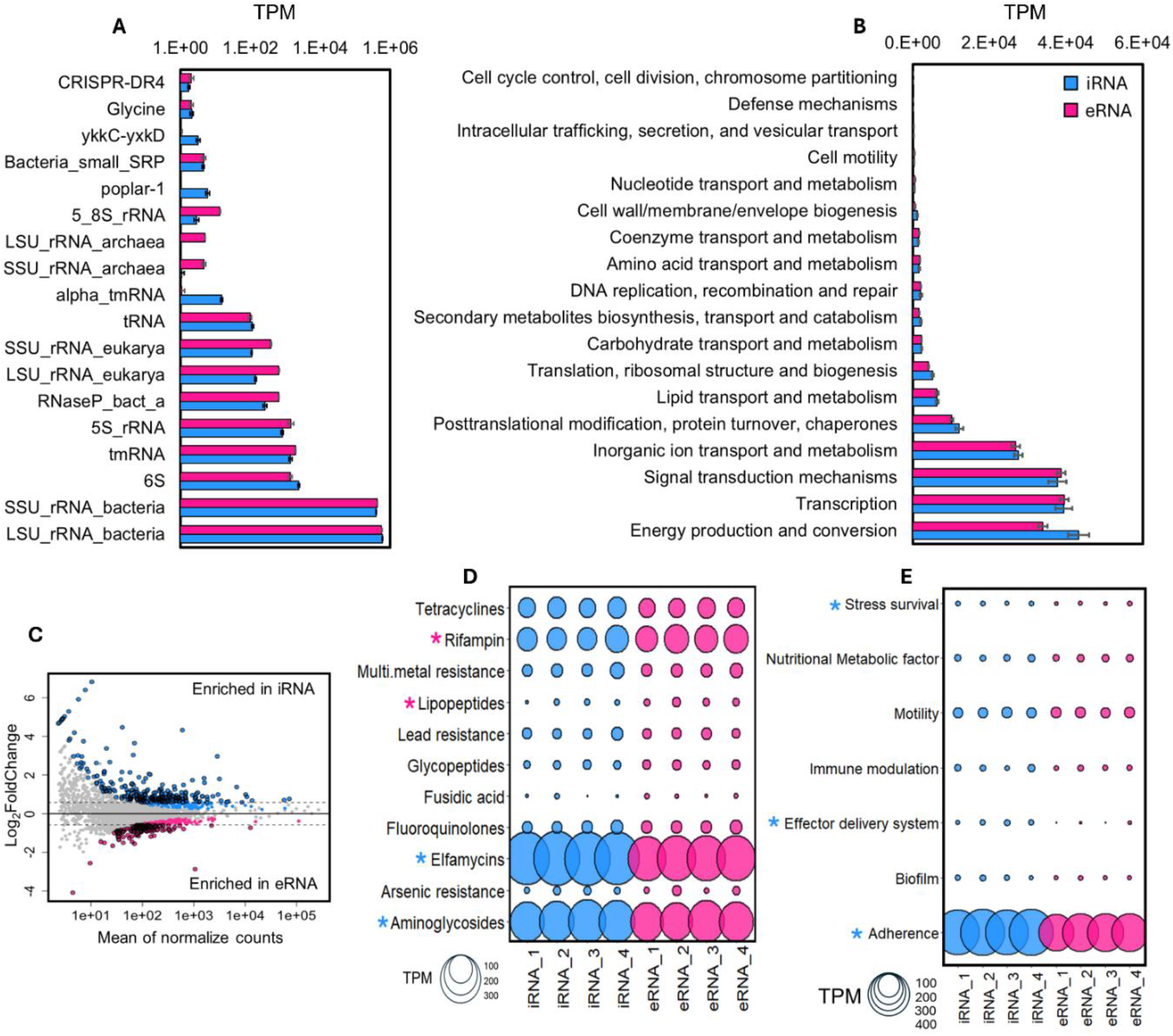
Features of drinking water metatranscriptome. **A)** Different types of RNA detected in iRNA and eRNA based on the Rfam database **B)** General functions expressed in the drinking water metatranscriptome by iRNA and eRNA according to COG categories. **C)** Fold change of different COGs. Significantly different COGs defined as COGs with p-adjusted < 0.01 and |log_2_(fold-change)| > 0.58. **D)** Antibiotic, biocide and metal resistance genes and **E)** Virulence factors expressed in the drinking water metatranscriptome based on metagenomic annotations from MEGARes^15^ and Virulence Factor database^16^. Statistically significant difference for **D)** and **E)** is indicated by ‘**\***’ (p < 0.05). ‘**\***’ indicates significant enrichment in iRNA and ‘**\***’ indicates significant enrichment in eRNA TPM: Transcripts per million.

Messenger RNA (mRNA) was recovered from both eRNA and iRNA (Figure 4B). The most prominent functions detected include energy production and conversion (such as nitrification, carbon metabolism, and ATP synthesis), transcription (DNA-binding response regulators, DNA-directed RNA polymerase, and cold shock proteins), and signal transduction mechanisms (including histidine kinases, anti-sigma factors, and chemotaxis). The potential of eRNA as a reference in differential metatranscriptomics was assessed using 2,495 COGs with over 20 counts across all samples for differential abundance analysis. A total of 596 COGs were found to be differentially abundant between iRNA and eRNA (p-adjusted < 0.01). In terms of biological relevance, 253 and 145 COGs were significantly enriched in iRNA and eRNA, respectively (p-adjusted < 0.01 and |log_2_(fold-change)| > 0.58, Figure 4C).

According to both COG and KEGG annotations (Tables S7 and S8), the top discovered transcripts revealed that genes involved in nitrification (*pmoA-amoA, pmoB-amoB, pmoC-amoC, hao and nirK*) were highly expressed in the sampled DWDS, facilitating the conversion of ammonia to nitrite. Nitrite could be subsequently converted to nitric oxide by nitrite reductase, which is plausible given the use of MCA as the disinfectant^20^. Additionally, aa3-type cytochrome c-oxidase (*coxABC*), which is more efficient than other c-type oxidases in a nutrient limiting environments^21^), was found to be active in the DWDS. These functions were enriched in the iRNA fraction, underscoring their significance for survival in the sampled DWDS.

Transcripts of genes related to aminoglycoside-, and elfamycin-. resistance showed significantly higher abundance in the iRNA fraction, suggesting selection pressure exerted by these antibiotics (Figure 4D). Additionally, the transcripts of genes for adherence (*Ef-Tu, groEL*), effector delivery systems, and stress tolerance were also enriched in the iRNA fraction, although the total number of assigned transcripts was low across all categories, except for adherence and motility (Figure 4E).

### Pathway fluxes in the context of cellular decisions within the DWDS

To examine cellular decisions within the DWDS, KEGG annotations were employed. Gene Set Enrichment Analysis (GSEA) was conducted to identify significantly enriched pathways represented by expressed KOs (Figure 5A and Figure S7). Based on the fold-change values of the genes in these pathways (Figure 5B), each pathway included several genes that were enriched in both iRNA and eRNA. Assuming that RNA release is predominantly a result of cell permeabilization or lysis due to disinfectant stress, the genes enriched in iRNA would indicate cellular decisions and pathways that support cell survival within the DWDS.

**Figure 5.**
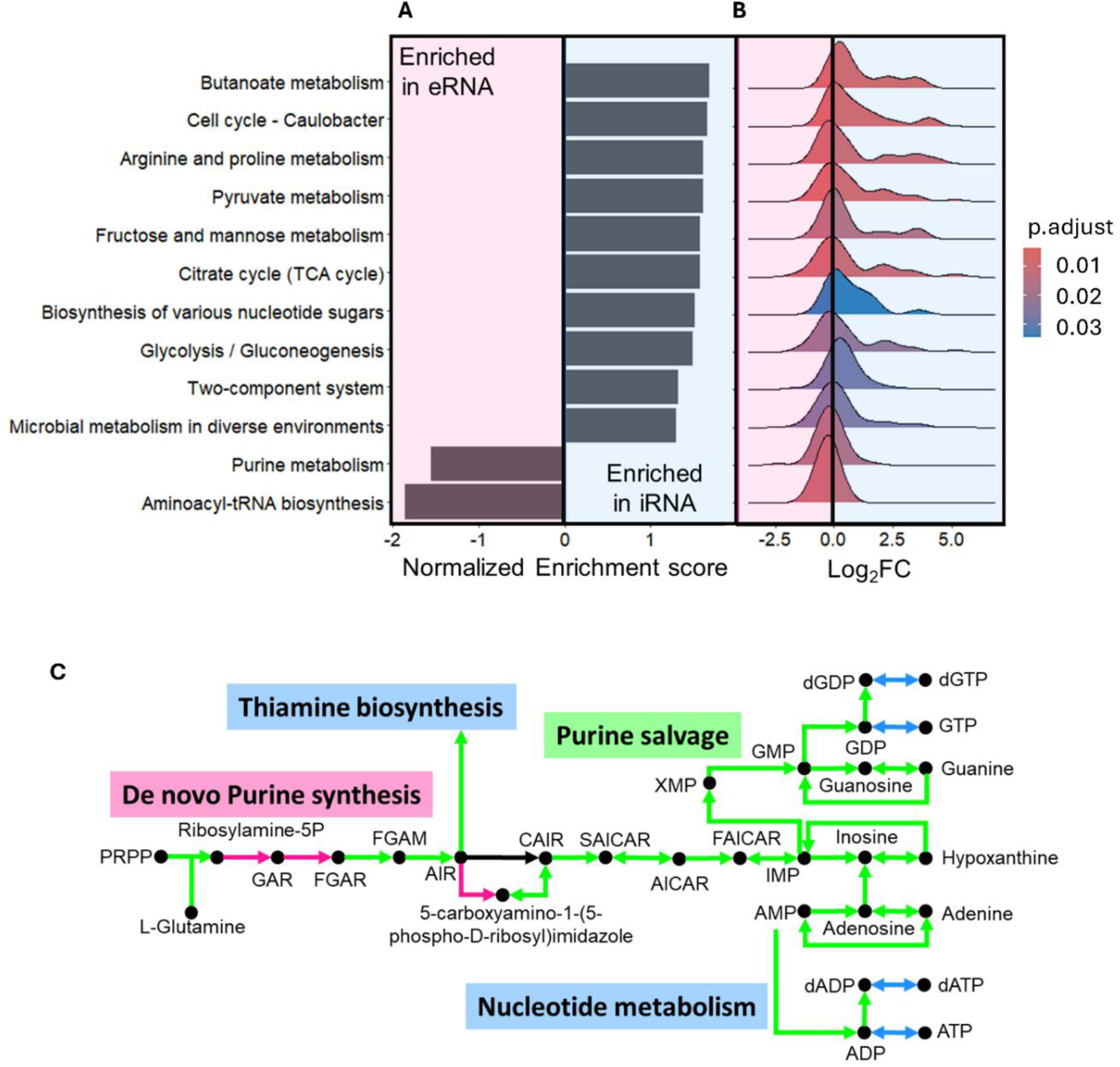
Gene set enrichment analysis on the significantly enriched KO groups (p-adjusted < 0.05 and log2foldchange > ± 0.58) with de novo purine synthesis as an example. **A)** Enrichment plot based on the normalized enrichment score showing enriched pathways in the iRNA and eRNA. **B)** Ridge plot showing the log_2_foldchange (Log_2_FC) of all genes involved in the enriched pathways. **C)** Reconstruction of the de novo purine synthesis pathway showing upregulated enzymes for substrate conversions within the pathway. Green represents enzymes/pathways not significantly enriched in iRNA and eRNA, light blue represents enzymes/pathways significantly enriched in iRNA while pink represents enzymes/pathways significantly enriched in eRNA (with p-adjusted < 0.01 and |log_2_(fold-change)| > 0.58). **Black arrows** represent enzymes/pathways **not detected** in the current data. PRPP: 5-Phosphoribosyl diphosphate; GAR: 5’-Phosphoribosylglycinamide; FGAR: N-Formyl-GAR; FGAM: 5’-Phosphoribosylformylglycinamidine; AIR: Aminoimidazole; CAIR: 1-(5’-Phosphoribosyl)-5-amino-4-imidazolecarboxylate; SAICAR: 1-(5’-Phosphoribosyl)-5-amino-4-(N-succinocarboxamide)-imidazole; AICAR: 5’-Phosphoribosyl-5-amino-4-imidazolecarboxamide; FAICAR: 5’-Phosphoribosyl-5-formamido-4-imidazolecarboxamide; IMP: Inosine monophosphate; GMP: Guanosine monophosphate; GDP: Guanosine diphosphate; dGDP: 2’-Deoxyguanosine 5’-diphosphate; GTP: Guanosine triphosphate; AMP: Adenosine monophosphate; ADP: Adenosine diphosphate; dADP: 2’-Deoxyadenosine 5’-diphosphate; ATP: Adenosine triphosphate; XMP: Xylulose monophosphate.

#### Energy efficiency, generation, and membrane regulation

To understand the differential pathways, it is essential to consider the context of the system. In an oligotrophic environment like the DWDS, energy efficiency is paramount. For instance, in purine metabolism (Figure 5C and S8), the de novo purine synthesis module shows multiple genes enriched in the eRNA fraction. This result suggests that cells relying on this module for purine synthesis might not thrive in the DWDS; instead, they utilized the more energy-efficient purine salvage pathway to recycle purine bases for DNA synthesis (dNTP) and generate energy-rich NTPs (Figure S9). The intermediate compound AIR (aminoimidazole) was channeled into thiamine synthesis, with relevant genes enriched in the iRNA (Figure S10). Additionally, enzymes responsible for dNTP/NTP formation were also enriched in the iRNA fraction (Figure S9). Furthermore, the process of energy producing fatty acid degradation was favored over fatty acid biosynthesis, which consumes energy (Figures S11–S13). Similar results for regulating cell membrane structure (Text S3, Figures S11, S14–16), carbon metabolism, and energy generation (Text S4. Figures S11C, S17-S18) can be found in the SI.

#### Synthesizing important cofactors

In the sampled DWDS, where MCA is used as a secondary disinfectant, genes associated with nitrification and assimilatory sulfate reduction were enriched in the iRNA (Figures 6A, B, S19, and S20). Siroheme is required as a prosthetic group for both nitrite and sulfite reductases, while heme is required for enzymes in the electron transfer chain^22^. Genes involved in the synthesis of both siroheme and heme, along with a heme transporter required for cytochrome biogenesis (ccmABCD, Figure 6C, and S21)^23^, were enriched in iRNA. In contrast, genes for the synthesis of the vitamin B12 coenzyme were enriched in eRNA, indicating a prioritization of heme synthesis as inhibiting heme synthesis promotes vitamin B12 production^24^. Furthermore, genes related to oxidative phosphorylation (Figure S22) and ribosome biogenesis (Figure S23) were also enriched in iRNA, reflecting the metabolic activity of cells thriving in the DWDS.

**Figure 6.**
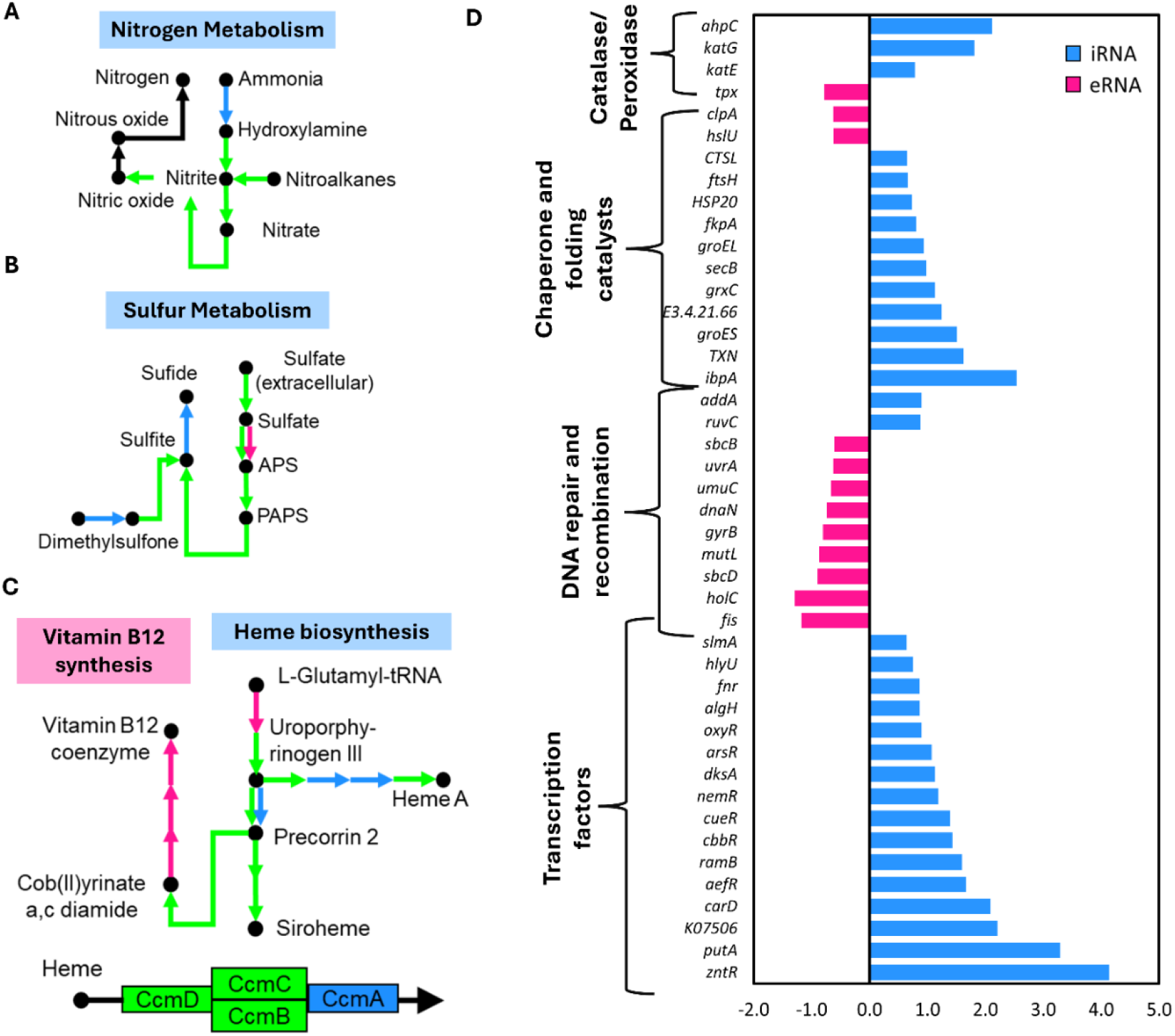
Factors important for the survival of cells in the presence of MCA as a disinfectant. **A)** Nitrogen metabolism in DWDS. Genes for nitrification and denitrification are enriched in the iRNA fraction. **B)** Sulfur metabolism in DWDS. Assimilatory sulfate reduction shows that sulfate can be used as an alternative electron acceptor in the DWDS. **C)** Synthesis of heme and siroheme as prosthetic groups is important for the survival of cells in the DWDS while synthesis of Vitamin B12 is reduced to channel flux into heme and siroheme synthesis **D)** Log_2_FoldChange of different transcription factors, chaperones, DNA repair enzymes and catalase/peroxidase which are significantly enriched in the either iRNA or eRNA fraction of the metatranscriptome. Green represents enzymes/pathways not significantly enriched in iRNA and eRNA, light blue represents enzymes/pathways significantly enriched in iRNA while pink represents enzymes/pathways significantly enriched in eRNA (with p-adjusted < 0.01 and |log_2_(fold-change)| > 0.58). **Black arrows** represent enzymes/pathways **not detected** in the current data. APS: Adenosine 5’-phosphosulfate; PAPS: 3’-Phosphoadenosine 5’-phosphosulfate.

#### Protein folding and stress response

Analysis of the BRITE categories from KEGG annotations (Figure S24) reveals that several DNA repair genes were enriched in eRNA (*uvrA, mutL, dnaN, holC, sbcD, sbcB, gyrB, fis and umuC*), suggesting DNA damage as a plausible cause of cell lysis (Figure 6D). In contrast, transcription factors were primarily enriched in iRNA (Figure 6D), helping cells adapt to the DWDS environment. For instance, the transcription factor *nemR* regulates detoxification in the presence of reactive chlorine species like MCA^25^, while *oxyR* controls the oxidative stress response induced by hydrogen peroxide^26^. Additionally, disulfide bond-reducing proteins (activators of the oxyR regulon) such as *grxC* was enriched in iRNA. The presence of oxidative stress due to MCA is further supported by the enrichment of catalases and peroxidase enzymes, such as *katE, katG*, and *ahpC*, in the iRNA fraction, indicating cellular responses to the disinfectant^27^. MCA can interact with proteins and inhibit their functions^28^. We found multiple chaperones responsible for the optimal folding of proteins and refolding of misfolded proteins (*hsp20, ibpA, fkpA, GroEL, GroES, atpB*), along with proteins responsible for turnover and quality control (*cathepsin L, ftsH, secB*) were enriched in iRNA. In contrast, ATP-dependent chaperones like *clpA* and *hslU* were found to be enriched in eRNA, highlighting the importance of chaperones for cell survival in the DWDS without compromising ATP resources.

The ability to obtain nutrients from decaying biomass components (amino acids, fatty acids etc.) is also essential for survival in oligotrophic DWDS^3,29^. In the current data set, degradation of proline to glutamate by putA is highly enriched in iRNA fraction (Figure 6D, S25). Glutamate can be used as a carbon source though the GABA shunt or glutamate dehydrogenase systems, though we did not capture expression of the whole pathway. Proline degradation might serve multiple purposes as it is shown to enhance oxidative stress tolerance in bacterial cells by enhancing *KatG* expression^30^ and methylglyoxal detoxification^31^.

## Discussion

Previously, environmental metatranscriptomes could only provide a general overview of microbial processes since differential analysis was not possible. This study demonstrates the use of eRNA to facilitate differential metatranscriptomics in disinfected drinking water systems. We identified active microorganisms (Figure 2D) and the factors that help these organisms to thrive in an oligotrophic environment even in the presence of a disinfectant (Figure 4–6, S11, S24).

Drinking water systems have been extensively studied using DNA-based methods, but the significance of eDNA in these systems has only recently been recognized^5^. While eDNA does not interfere with qualitative analysis of DWDS metagenomes, it obscures the quantitative analysis. The metagenome reveals a wide range of transposons and ARGs (Figure 3) in the DWDS, with approximately 50% being extracellular. Consequently, prior studies showing enhanced horizontal gene transfer through transformation during disinfection treatment^32,33^ are highly relevant in the context of actual DWDS environments. Additionally, we found the GC content of coding regions to greatly enhance persistence in the extracellular form. Future eARG studies should account for this factor while examining their persistence and fate in environmental matrices.

DNA–based studies in disinfected drinking water have reported the enrichment of aminoglycoside–, multidrug–, fosfomycin–, sulfonamide–, and bacitracin resistance genes post chlorination or chloramination^34–36^. In this study, we confirmed the presence of these resistance genes in addition to the high abundance of heavy metal resistance genes. However, most of the ARGs were enriched in the eDNA fraction, raising concerns on the accuracy of total DNA–based quantification of ARGs in drinking water, especially in disinfected systems. The metatranscriptome analyses of the DWDS revealed a high expression of ARGs for aminoglycosides and elfamycin resistance, which were enriched in iRNA (Figure 4D). This ARG expression aligns with pathway analysis. Aminoglycosides and elfamycin inhibit polypeptide synthesis^37^ by preventing transfer RNAs (tRNAs) from accessing the active site. In pathway analysis (Figure 5A, S26), aminoacyl-tRNA was enriched in eRNA, suggesting selection pressure exerted by antimicrobials and disinfectants in the DWDS.

The metatranscriptome data obtained in this study can be used to validate conclusions from drinking water metagenomic studies. For instance, based on metagenome analysis, the pathways for oxidation of methane to methanol, fatty acid metabolism, TCA cycle and glycosyltransferase mediated modification of polysaccharides are key to survival in the DWDS^38^. Potential to utilize microbial decay products through glyoxylate shunt and propanoyl–CoA metabolism are additional factors for success in disinfected drinking water systems^3^. Our metatranscriptome data shows expression of all these pathways, with multiple steps enriched in the iRNA fraction (Figure S11). Additionally, we show that energy efficiency plays a crucial role in the success of cells in the DWDS, with an emphasis on enhancing and maintaining flux into the TCA cycle, fatty acid degradation, and potential utilization of products from decaying biomass. Furthermore, Potgieter et al.^20^ suggested the reduction of nitrate to nitric oxide in chloraminated drinking water for biofilm regulation. Our results confirm the expression of this pathway in drinking water (Figure 6A). However, we did not detect expression of genes for further reduction of the nitric oxide. These metabolic pathways are dependent on synthesis of key prosthetic groups, such as heme, siroheme, and ubiquinone -essential for electron transfer and energy generation - are also a priority for organisms in the DWDS.

Metagenome assembled genomes (MAGs) from disinfected DWDS are associated with high SOS–response genes^3^ indicating importance of oxidative stress response genes for detoxification and the prevention of DNA damage. Different stress response mechanisms were identified using the metatranscriptomics data (Figure 6D). Maintaining a dynamic cell membrane structure through phospholipid transporters, chaperone-assisted protein folding, and the prevention of misfolded protein aggregation, along with recycling and quality control, is vital for survival in the oligotrophic DWDS.

The different RNA subtypes detected in eRNA (Figure 4A) such as tmRNA and sRNA, may have diverse implications for organism interactions beyond DWDS. Bacterial sRNA and noncoding RNA packed in outer membrane vesicles play an integral role in inter kingdom communication^39–42^. With RNA being persistent in the extracellular form, similar interactions may be common in microbial ecology, orchestrating microbiome interactions. For instance, the most abundant tmRNA, SsrA, is known to suppress immune and inflammatory responses in squids when interacting with the pathogenic *Vibrio fischeri*^43^. Similarly, another sRNA Xosr001 from *Xanthomonas oryzae* can suppress stomatal immunity in rice^44^.

We are just beginning to grasp the various aspects of eRNA, and this study represents a significant step forward in enhancing our understanding of microbial ecological interactions in both natural and engineered environments. This study analyses four biological replicates collected in parallel and presents an in-depth analysis of the four nucleic acid fractions and their importance. Future studies could involve multiple sampling locations and time points to thoroughly investigate disinfectant-bacteria interactions and metabolism across geographically and operationally distinct drinking water systems. Lastly. it demonstrates the utility of eRNA as a reference to study in-situ bacterial responses under defined stress conditions.

## Methods

### Drinking water sampling and experimental design

Drinking water was collected directly from the tap using glass fiber membranes (Advantec, 0.4 µm nominal pore size, 25 mm diameter) housed in polypropylene in-line filter holders (Sterlitech), connected to a 4-way flow splitter. The tap was opened and allowed to flush for 5 minutes before the sampling apparatus was attached to the tap. The glass fiber membrane was chosen as i) it can effectively retain more eDNA as compared to other filter membranes^45^, ii) frequently used for water sampling and environmental DNA extraction^46^, and iii) provides high flowthrough enabling direct sampling from the tap. While the glass fibre membrane does not directly bind eRNA, its high flowthrough property is essential to capture high amount of dead biomass on the filter membrane. More details about the sampling method and distribution system can be found in our previous work^5^. The initial flow rate ranged from 300 to 315 mL/min, and filtration continued until the filters became clogged, processing a total of ∼20 L of tap water for each filter. The flow rate was monitored every 30 minutes to calculate the volume of water filtered. For sequencing, the sampling continued with a fresh set of filters after the first set was clogged (Figure 1A). All collected samples were processed immediately. Further details on sample processing are included in the DNA extraction sections.

Water samples were collected at the start (time 0), middle (time 2), and end (time 4) of the sampling process. 1 mL of the collected water samples was stained with 1 µl each of SYTO9 and Propidium Iodide. A total of 700 µl of the stained drinking water was analysed using a BD Accuri C6 flow-cytometer, with a threshold of 1000 on FL1-H to count the intact cells and assess the phenotypic biodiversity of the water samples at each sampling stage using the ‘Phenoflow’ package. Unstained water samples were used as controls to exclude any spurious signals^11,12^.

Finished water quality parameters such as monochloramine (HR TNT, Method 10172), nitrate (TNT 835), nitrite (TNT 839), ammonia (TNT 830), and free chlorine (TNT 866) were measured using the respective HACH TNT kits. TOC analyzer was used to measure the total carbon and nitrogen. The detailed water chemistry is provided in Supporting Information (SI) Table S2. The water quality parameters were measured at the end of the sampling process.

### eNAC extraction

After sampling, the clogged filters were collected and processed immediately. One half of each filter was used to extract iDNA and eDNA, while the other half was used for iRNA and eRNA (Figure 1A).

eNAC was extracted from the filters following previous protocols^47,48^ (Figure 1C). Briefly, the filters were first treated with 1 mg/mL Proteinase K (c). The P_k_ treatment facilitates the breakdown of cell aggregates and release of DNA/RNA associated with proteins while TE buffer helps to solubilize the extracellular DNA/RNA. P_k_ treatment does not affect cell viability and permeability (Additional information in SI). Following the P_k_ treatment, eDNA and eRNA were recovered from the filters. For eDNA extraction, 3 washes using NaP buffer were performed. The supernatant from the wash was centrifuged to collect washed off cells (subsequently included in iNAC extraction) and the supernatant was used to purify eDNA. For eRNA samples, washing with NaP did not result in recovery of additional eRNA. Thus, eRNA was recovered only from the initial TE buffer+ P_k_ treatment.

For sequencing, the supernatants from 2 filter sets were pooled at this point for each sample. Pooling was done from the same filter holder position in the continuous sampling design. The eNAC was precipitated overnight at 4 °C using 1/10^th^ volume of 5 M sodium acetate (pH 5) and 1 volume of isopropanol. The samples were then centrifuged at 12,000 × g for 15 min to pellet the eNAC, washed with 70% ethanol, and resuspended in RNase/DNase-free water. The samples were treated with DNase/RNase and were subsequently purified using the appropriate Zymogen clean and concentrator kit^10^. All extraction buffers were freshly prepared before each extraction, filter sterilized and subjected to UV treatment as an additional precaution. For eDNA and eRNA extraction, the sample buffers served as blanks, with no nucleic acids detected (sample concentration < 0.4 ng/µL for RNA and <0.01 ng/µL for DNA). Moreover, the eNAC and iNAC samples were processed using separate buffers for extraction, minimizing the risk of cross-contamination between these buffers.

### iNAC extraction

After extracting eNAC, the respective filters and cell pellets (obtained after centrifugation of eNAC extracts) were pooled for iNAC extraction. iDNA was extracted following our previously published protocol^5^. Briefly, samples were homogenized in 2 mL Eppendorf tubes containing 1 mL of pre-warmed CTAB-proteinase K buffer (60 °C), 0.1 g of acid-washed glass beads (400-600 nm in diameter), and a larger glass bead with a 2 mm radius, using a Genie 2 vortex machine (Scientific Industries) at maximum speed (3,200 rpm) for 10 min. The samples were then incubated at 60 °C for 1 h and centrifuged for 5 min at 10,000 × g to remove debris. The supernatant was collected, and 1 mL of chloroform/isoamyl alcohol (v/v 24:1) was added. The samples were mixed by inversion for 2 min at room temperature, followed by centrifugation at 14,000 × g and 4 °C for 10 min. The aqueous phase was collected and treated with 5 µL of RNase A (20 mg/mL, Invitrogen) for 30 min at 37 °C. One volume of isopropanol was added, and the samples were incubated at room temperature overnight, then centrifuged at 14,000 × g and 4 °C for 15 min. The DNA pellet was washed twice with 70% ethanol, and a final centrifugation was carried out at 21,130 × g for 15 min. The pellet was air-dried and resuspended in nuclease free water. DNA was purified using the Zymogen DNA Clean and Concentrator Kit.

iRNA extraction was initiated immediately after the P_k_ treatment. This results in a time lag of 30 mins between sampling and RNA extraction. Metatranscriptomics in low density systems is still a challenge, and we cannot completely rule out some alteration in the transcriptome profile during sample collection and processing. However, these effects may be considered minimal given the quick sample processing time and comparative nature of our study^49^. Use of reagents like RNAlater or similar kind were avoided to preserve the distinction between intracellular and extracellular fractions.

iRNA was extracted using the Trizol method. Briefly, immediately after the P_k_ treatment, the filter was homogenized in 1.5 mL of Trizol reagent, along with 0.1 g of acid-washed glass beads (400-600 nm in diameter) and a larger glass bead (2 mm radius), using the Genie 2 vortex machine (Scientific Industries) at the maximum speed (3200 rpm) for 10 min. The samples were incubated at 60 °C for 10 min before returning to room temperature. After adding 300 µL of chloroform, the samples were gently mixed by inversion and incubated for 5 min at room temperature. Centrifugation at 12,000 g for 10 min facilitated the formation of the aqueous phase, which was transferred to a fresh tube and extracted with chloroform/isoamyl alcohol (24:1 v/v) to remove residual phenol from the Trizol. The aqueous phase was mixed with one volume of isopropanol and incubated overnight at 4 °C. The samples were then centrifuged at 16,000 g for 30 min to precipitate the iRNA, washed with 70% ethanol, and resuspended in pre-warmed nuclease-free water. The samples were treated with DNase and purified using the Zymogen RNA Clean and Concentrator Kit. The quality of the extracted nucleic acids was assessed using a Nanodrop, and quantification was performed using the appropriate Qubit kits.

Extraction from filter blanks were also carried out (n = 4) and the nucleic acid were measured using their respective Qubit high sensitivity kits. Nucleic acids remained undetectable in all extracts (sample concentration < 0.4 ng/µL for RNA and <0.01 ng/µL for DNA).

DNA samples were sequenced using MiSeq 300bp paired-end sequencing. The TrueSeq stranded library preparation kit was used to process RNA samples following the low sample protocol without mRNA purification, with 200 ng of total RNA as template. The first strand of cDNA was synthesized using SuperScript II reverse transcriptase enzyme, followed by second strand synthesis to produce double-stranded (ds) cDNA with blunt ends. The library was sequenced using HiSeq 100bp paired-end sequencing at the SCELSE sequencing facility in Singapore that specialises in sequencing low biomass samples. Four replicates of each NAC type (eDNA, iDNA, eRNA, and iRNA) were sequenced. Raw sequences have been deposited in NCBI-SRA under the bioproject PRJNA1036395.

To prevent kit/reagent–based contamination, following precautions were taken: i) The sequencer is always washed using tween 20% and milli-Q water between sequencing runs, ii) Flow cells are individually packed, no cross-contamination between flow cells, iii) Libraries were sequenced in a dedicated flow cell, with no other samples multiplexed together with our samples, and iv) No clinical samples were run on the machines prior to our run. Furthermore, DNA and RNA were sequenced on separate flow cells and library preparation was executed using different kits. High alignment rates (>99.7 % alignment of RNA sequences on DNA assembly) further indicate high quality sequencing performance.

### Sequencing analysis for community determination

The quality of the raw reads was assessed using FastQC^50^. Adapters and low quality reads were removed using Trimmomatic (LEADING:8 TRAILING:8 SLIDINGWINDOW:10:15 MINLEN:30)^51^. Taxonomic classification of the samples was performed with Kraken2^52^, utilizing the Kraken2 standard database (release 9/4/2024) with --minimum-hit-groups value of 3 (default is 2). The output was filtered and visualized using Pavian^53^ to include only assignments at the genus level, which was used for subsequent analyses in R (v4.4) using the “phyloseq”^54^ and “vegan”^55^ packages. Plots were generated using the package “ggplot2”^56^. DNA and RNA samples were processed separately due to the differences in library sizes. The taxa table was rarefied to the lowest number of sequences obtained for any sample in the dataset (DNA = 852,146 and RNA = 25,911,012) using the *rarefy_even_depth()* function followed by removal of taxa with less than 10 counts across all samples. The α-diversity indices, such as “Observed” and “Shannon”, were calculated for the eNAC and iNAC samples. Principal Co-ordinate Plots (PCoA) were constructed based on the Bray-Curtis dissimilarity matrix, which was calculated from the rarefied dataset. Hierarchical cluster analysis was conducted using the average linkage clustering method^57^. Non parametric β-diversity analyses were conducted using the *adonis2()*^58^, *betadisper()*^59^, and *permutest()* functions from the “vegan” package, to assess variance among samples. Statistical significance of all datasets was determined using pairwise Student’s t-tests for means or Analysis of Variance (ANOVA) in Excel. The fold-change in detection of a genera between iNAC and eNAC was calculated using the DeSeq2.

### Metagenome and metatranscriptome analysis

The quality-controlled reads were processed using the SqueezeMeta pipeline^60^ in co-assembly mode with Megahit^61^ as the assembler. This automated pipeline facilitates metagenomic and metatranscriptomic analyses, generating all necessary outputs, including alignment files, DNA assembly, predicted coding regions, amino acid sequences, KEGG, COG, and EggNOG annotations, raw counts, GC content of predicted coding sequences, abundance matrices, transcripts per million (TPM) values, along with COG and KO counts. RNA sequences were aligned to the assembled metagenome to assess the expression of various coding regions. TPM values were recalculated after removing rRNA-aligned reads using SQMtools^62^.

The assembled DNA sequences were aligned to the Rfam^18^ database using Infernal^17,63^ to identify different types of RNA present in the assembly. BAM files generated by the pipeline, which show the alignment of RNA to the metagenome assembly, were) analyzed with featureCount^64^ to quantify the RNA assigned to each subtype. The MEGARes database^15^ and Virulence Factor database^16^ were used to align the predicted protein sequences using DIAMOND^65^. Annotated sequences were filtered based on an e-value (< 10^-5^) and sequence identity (> 70%) to identify antimicrobial and heavy metal resistance genes (ARGs) and virulence factors (VFs) in the metagenome. TPM values for the annotated ARGs and VFs were extracted from the SqueezeMeta output (*. mapcount file). Data were aggregated according to different classes of antibiotics and virulence genes, and Student’s t-test for means was used to assess significantly enriched categories in iNAC and eNAC.

Raw counts for KOs (> 5) and COGs (> 20) were filtered for count and analyzed using DeSeq2^66^ to determine differential abundance of various functional groups. Differentially abundant COGs were selected based on a criterion of p-adjusted < 0.01 and |log_2_(fold-change)| > 0.58, while differentially abundant KOs were filtered with p-adjusted < 0.1 and |log_2_(fold-change)| > 0.58. The KOs were further analyzed using the “*Clusterprofiler*” package in R^67^ for gene set enrichment analysis. Differentially abundant KOs were manually analyzed using KEGG color and KEGG mapper tools^68^.

## Associated content

### Supporting Information

Additional information on flow cytometry based cell count and phenotypic fingerprinting (Fig. S1), eNAC extraction and proteinase K treatment (Fig. S2) α– diversity, clustering, and dispersion of microbial community (Fig. S3), comparison of microbial community detection using eDNA, iDNA, eRNA and iRNA (Fig. S4), drinking water community composition (Fig S5, S6), enriched and depleted metabolic pathways in iRNA and eRNA (Fig S7), KEGG metabolic pathways for transcriptome analyses (Fig. S8–S26), water quality parameters (Table S1), nucleic acid concentrations (Table S2), read classification statistics by Kraken and Pavian (Table S3), β–diversity statistics (Table S4), top 30 COGs and KOs in DNA (Tables S5 and S6),), top 30 COGs and KOs in RNA (Tables S5 and S6) are provided in the supplementary file.

## Acknowledgements

This research was supported by the Ministry of Education Singapore (MOE AcRF Tier 1 grant, Award No.: RT10/22) and by the MOE and NRF and under its Research Centre of Excellence Programme, Singapore Centre for Environmental Life Sciences Engineering (M4330005.C70 to B.C.), Nanyang Technological University, Singapore.

## Author Contributions

SB: Writing – review & editing, Writing – original draft, Visualization, Methodology, Investigation, Formal analysis, Data curation, Conceptualization; SM Writing – original draft, Methodology, Investigation; DC: Writing – review & editing, Methodology, Investigation, Formal analysis, Data curation; YL: Writing – review & editing, Investigation, Formal analysis, SK: Writing – review & editing, Supervision, Resources, Methodology; BC: Writing – review & editing, Supervision, Resources, Methodology, Funding acquisition, Conceptualization.

## Competing Interest Statement

The authors declare no competing financial interests.

## Notes

### Competing Interest Statement

The authors have declared no competing interest.

## REFERENCES

1. Hull, N. M. et al. Drinking Water Microbiome Project: Is it Time? Trends in Microbiology 27, 670–677 (2019).

2. Waak, M. B., Hozalski, R. M., Hallé, C. & LaPara, T. M. Comparison of the microbiomes of two drinking water distribution systems—with and without residual chloramine disinfection. Microbiome 7, 87 (2019).

3. Dai, Z. et al. Disinfection exhibits systematic impacts on the drinking water microbiome. Microbiome 8, 42 (2020).

4. Abkar, L., Moghaddam, H. S. & Fowler, S. J. Microbial ecology of drinking water from source to tap. Science of The Total Environment 908, 168077 (2024).

5. Sakcham, B., Kumar, A. & Cao, B. Extracellular DNA in Monochloraminated Drinking Water and Its Influence on DNA-Based Profiling of a Microbial Community. Environ. Sci. Technol. Lett. 6, 306–312 (2019).

6. Inkinen, J. et al. Diversity of ribosomal 16S DNA- and RNA-based bacterial community in an office building drinking water system. Journal of Applied Microbiology 120, 1723– 1738 (2016).

7. Li, R. et al. Comparison of DNA-, PMA-, and RNA-based 16S rRNA Illumina sequencing for detection of live bacteria in water. Sci Rep 7, 5752–5752 (2017).

8. Zhang, Y. & Liu, W.-T. The application of molecular tools to study the drinking water microbiome – Current understanding and future needs. Critical Reviews in Environmental Science and Technology 49, 1188–1235 (2019).

9. Abu-Ali, G. S. et al. Metatranscriptome of human faecal microbial communities in a cohort of adult men. Nat Microbiol 3, 356–366 (2018).

10. Bairoliya, S., Goel, A., Mukherjee, M., Koh Zhi Xiang, J. & Cao, B. Monochloramine Induces Release of DNA and RNA from Bacterial Cells: Quantification, Sequencing Analyses, and Implications. Environ. Sci. Technol. 56, 15791–15804 (2022).

11. Favere, J., Buysschaert, B., Boon, N. & De Gusseme, B. Online microbial fingerprinting for quality management of drinking water: Full-scale event detection. Water Research 170, 115353 (2020).

12. Props, R., Monsieurs, P., Mysara, M., Clement, L. & Boon, N. Measuring the biodiversity of microbial communities by flow cytometry. Methods in Ecology and Evolution 7, 1376–1385 (2016).

13. Aggarwal, S. et al. Effects of Chloramine and Coupon Material on Biofilm Abundance and Community Composition in Bench-Scale Simulated Water Distribution Systems and Comparison with Full-Scale Water Mains. Environmental Science & Technology 52, 13077–13088 (2018).

14. Chiao, T.-H., Clancy, T. M., Pinto, A., Xi, C. & Raskin, L. Differential Resistance of Drinking Water Bacterial Populations to Monochloramine Disinfection. Environmental Science & Technology 48, 4038–4047 (2014).

15. Bonin, N. et al. MEGARes and AMR++, v3.0: an updated comprehensive database of antimicrobial resistance determinants and an improved software pipeline for classification using high-throughput sequencing. Nucleic Acids Research 51, D744–D752 (2023).

16. Liu, B., Zheng, D., Zhou, S., Chen, L. & Yang, J. VFDB 2022: a general classification scheme for bacterial virulence factors. Nucleic Acids Research 50, D912–D917 (2021).

17. Nawrocki, E. P. & Eddy, S. R. Infernal 1.1: 100-fold faster RNA homology searches. Bioinformatics 29, 2933–2935 (2013).

18. Kalvari, I. et al. Rfam 14: expanded coverage of metagenomic, viral and microRNA families. Nucleic Acids Research 49, D192–D200 (2020).

19. Ghosal, A. et al. The extracellular RNA complement of Escherichia coli. MicrobiologyOpen 4, 252–266 (2015).

20. Potgieter, S. C., Dai, Z., Venter, S. N., Sigudu, M. & Pinto, A. J. Microbial Nitrogen Metabolism in Chloraminated Drinking Water Reservoirs. mSphere 5, e00274–20 (2020).

21. Wu, Y., Zaiden, N., Liu, X., Mukherjee, M. & Cao, B. Responses of Exogenous Bacteria to Soluble Extracellular Polymeric Substances in Wastewater: A Mechanistic Study and Implications on Bioaugmentation. Environmental Science & Technology 54, 6919–6928 (2020).

22. Crane, B. R. & Getzoff, E. D. The relationship between structure and function for the sulfite reductases. Current Opinion in Structural Biology 6, 744–756 (1996).

23. Thöny-Meyer, L., Fischer, F., Künzler, P., Ritz, D. & Hennecke, H. Escherichia coli genes required for cytochrome c maturation. Journal of Bacteriology 177, 4321–4326 (1995).

24. Biedendieck, R. et al. Metabolic engineering of cobalamin (vitamin B12) production in Bacillus megaterium. Microbial Biotechnology 3, 24–37 (2010).

25. Gray, M. J., Wholey, W.-Y., Parker, B. W., Kim, M. & Jakob, U. NemR Is a Bleach-sensing Transcription Factor. Journal of Biological Chemistry 288, 13789–13798 (2013).

26. Holder, D., Berry, D., Dai, D., Raskin, L. & Xi, C. A dynamic and complex monochloramine stress response in Escherichia coli revealed by transcriptome analysis. Water Research 47, 4978–4985 (2013).

27. Wan, F., Yin, J., Sun, W. & Gao, H. Oxidized OxyR Up-Regulates ahpCF Expression to Suppress Plating Defects of oxyR- and Catalase-Deficient Strains. Frontiers in Microbiology 10, (2019).

28. Levine, R. L., Mosoni, L., Berlett, B. S. & Stadtman, E. R. Methionine residues as endogenous antioxidants in proteins. Proceedings of the National Academy of Sciences 93, 15036–15040 (1996).

29. Sudarshan, A. S. et al. New Drinking Water Genome Catalog Identifies a Globally Distributed Bacterial Genus Adapted to Disinfected Drinking Water Systems. Environ. Sci. Technol. 58, 16475–16487 (2024).

30. Zhang, L., Alfano, J. R. & Becker, D. F. Proline Metabolism Increases katG Expression and Oxidative Stress Resistance in Escherichia coli. Journal of Bacteriology 197, 431– 440 (2015).

31. Berney, M., Weimar, M. R., Heikal, A. & Cook, G. M. Regulation of proline metabolism in mycobacteria and its role in carbon metabolism under hypoxia. Molecular Microbiology 84, 664–681 (2012).

32. Zhang, S. et al. Chlorine disinfection facilitates natural transformation through ROS-mediated oxidative stress. The ISME Journal 15, 2969–2985 (2021).

33. Zhang, Y., Gu, A. Z., He, M., Li, D. & Chen, J. Subinhibitory Concentrations of Disinfectants Promote the Horizontal Transfer of Multidrug Resistance Genes within and across Genera. Environmental Science & Technology 51, 570–580 (2017).

34. Ma, L., Yang, H., Guan, L., Liu, X. & Zhang, T. Risks of antibiotic resistance genes and antimicrobial resistance under chlorination disinfection with public health concerns. Environment International 158, 106978 (2022).

35. Sevillano, M. et al. Differential prevalence and host-association of antimicrobial resistance traits in disinfected and non-disinfected drinking water systems. Science of The Total Environment 749, 141451 (2020).

36. Tiwari, A. et al. Bacterial Genes Encoding Resistance Against Antibiotics and Metals in Well-Maintained Drinking Water Distribution Systems in Finland. Front. Microbiol. 12, (2022).

37. Kotra, L. P., Haddad, J. & Mobashery, S. Aminoglycosides: Perspectives on Mechanisms of Action and Resistance and Strategies to Counter Resistance. Antimicrobial Agents and Chemotherapy 44, 3249–3256 (2000).

38. Liu, H. et al. Functional traits and health implications of the global household drinking-water microbiome retrieved using an integrative genome-centric approach. Water Research 250, 121094 (2024).

39. Pita, T., Feliciano, J. R. & Leitão, J. H. Extracellular RNAs in Bacterial Infections: From Emerging Key Players on Host-Pathogen Interactions to Exploitable Biomarkers and Therapeutic Targets. Int J Mol Sci 21, 9634 (2020).

40. Felden, B. & Augagneur, Y. Diversity and Versatility in Small RNA-Mediated Regulation in Bacterial Pathogens. Front. Microbiol. 12, (2021).

41. Grenier, T. et al. Intestinal GCN2 controls Drosophila systemic growth in response to Lactiplantibacillus plantarum symbiotic cues encoded by r/tRNA operons. Elife 12, e76584 (2023).

42. He, M. et al. Insights into the regulatory role of bacterial sncRNA and its extracellular delivery via OMVs. Appl Microbiol Biotechnol 108, 29 (2023).

43. Moriano-Gutierrez, S. et al. The noncoding small RNA SsrA is released by Vibrio fischeri and modulates critical host responses. PLOS Biology 18, e3000934 (2020).

44. Wu, Y. et al. Suppression of host plant defense by bacterial small RNAs packaged in outer membrane vesicles. Plant Comm 5, (2024).

45. Siuda, W. & Güde, H. Determination of dissolved deoxyribonucleic acid concentration in lake water. Aquatic Microbial Ecology 11, 193–202 (1996).

46. Eichmiller, J. J., Miller, L. M. & Sorensen, P. W. Optimizing techniques to capture and extract environmental DNA for detection and quantification of fish. Molecular Ecology Resources 16, 56–68 (2016).

47. Ogram, A., Sayler, G. S. & Barkay, T. The extraction and purification of microbial DNA from sediments. Journal of Microbiological Methods 7, 57–66 (1987).

48. Liu, M., Hata, A., Katayama, H. & Kasuga, I. Consecutive ultrafiltration and silica adsorption for recovery of extracellular antibiotic resistance genes from an urban river. Environmental Pollution 260, 114062 (2020).

49. Lesniewski, R. A., Jain, S., Anantharaman, K., Schloss, P. D. & Dick, G. J. The metatranscriptome of a deep-sea hydrothermal plume is dominated by water column methanotrophs and lithotrophs. ISME J 6, 2257–2268 (2012).

50. Andrews, S. FastQC A Quality Control tool for High Throughput Sequence Data.

51. Bolger, A. M., Lohse, M. & Usadel, B. Trimmomatic: a flexible trimmer for Illumina sequence data. Bioinformatics 30, 2114–20 (2014).

52. Wood, D. E., Lu, J. & Langmead, B. Improved metagenomic analysis with Kraken 2. Genome Biology 20, 257 (2019).

53. Breitwieser, F. P. & Salzberg, S. L. Pavian: interactive analysis of metagenomics data for microbiome studies and pathogen identification. Bioinformatics 36, 1303–1304 (2020).

54. McMurdie, P. J. & Holmes, S. phyloseq: An R Package for Reproducible Interactive Analysis and Graphics of Microbiome Census Data. PLOS ONE 8, e61217 (2013).

55. Dixon, P. VEGAN, a package of R functions for community ecology. Journal of Vegetation Science 14, 927–930 (2003).

56. Wickham, H. Ggplot2: Elegant Graphics for Data Analysis. (Springer Publishing Company, Incorporated, 2009).

57. Eisen, M. B., Spellman, P. T., Brown, P. O. & Botstein, D. Cluster analysis and display of genome-wide expression patterns. Proceedings of the National Academy of Sciences 95, 14863–14868 (1998).

58. J., A. M. A new method for non-parametric multivariate analysis of variance. Austral Ecology 26, 32–46 (2001).

59. J., A. M., E., E. K. & H., M. B. Multivariate dispersion as a measure of beta diversity. Ecology Letters 9, 683–693 (2006).

60. Tamames, J. & Puente-Sánchez, F. SqueezeMeta, A Highly Portable, Fully Automatic Metagenomic Analysis Pipeline. Frontiers in Microbiology 9, (2019).

61. Li, D., Liu, C.-M., Luo, R., Sadakane, K. & Lam, T.-W. MEGAHIT: an ultra-fast single-node solution for large and complex metagenomics assembly via succinct de Bruijn graph. Bioinformatics 31, 1674–1676 (2015).

62. Puente-Sánchez, F., García-García, N. & Tamames, J. SQMtools: automated processing and visual analysis of ‘omics data with R and anvi’o. BMC Bioinformatics 21, 358 (2020).

63. Kalvari, I. et al. Non-Coding RNA Analysis Using the Rfam Database. Current Protocols in Bioinformatics 62, e51 (2018).

64. Liao, Y., Smyth, G. K. & Shi, W. featureCounts: an efficient general purpose program for assigning sequence reads to genomic features. Bioinformatics 30, 923–930 (2013).

65. Buchfink, B., Xie, C. & Huson, D. H. Fast and sensitive protein alignment using DIAMOND. Nature Methods 12, 59–60 (2015).

66. Love, M. I., Huber, W. & Anders, S. Moderated estimation of fold change and dispersion for RNA-seq data with DESeq2. Genome Biology 15, 550 (2014).

67. Wu, T. et al. clusterProfiler 4.0: A universal enrichment tool for interpreting omics data. The Innovation 2, 100141 (2021).

68. Kanehisa, M., Furumichi, M., Sato, Y., Kawashima, M. & Ishiguro-Watanabe, M. KEGG for taxonomy-based analysis of pathways and genomes. Nucleic Acids Research 51, D587–D592 (2022).

