## Supplementary material for "From short-lived to persistent: The significance of extracellular RNA in disinfected drinking water microbiomes": SI

**This file includes the following:**

No. of Pages: 38

No. of Text: 4

No. of Figures: 26

No. of Tables: 8

**Text S1:**

**Proteinase K ($P_k$) treatment enhances eNAC extraction.**

$P_k$ treatment of the collected filters significantly increased the eNAC recovery (Figure S2A, B). 120 ± 5 ng of eDNA could be recovered from the control filters which increased to 217 ± 7 ng of eDNA after $P_k$ treatment. eRNA could be detected in only 3 out of the 4 samples without $P_k$ treatment. eRNA recovery increased from 340 ± 7 ng to 619 ± 168 ng after $P_k$ treatment. $P_k$ treatment does not affect cell viability and permeability (Figure S2C). It effectively liberates protein bound DNA and RNA by breaking cell aggregates corroborated by increased CFU detection (Figure S2C) and reduced average cell size (Figure S2D, E). After $P_k$ treatment, the subsequent extraction was carried out using NaP buffer.

**Text S2:**

To evaluate the functional potential of the DWDS microbiome, we co-assembled a total of 27,710,000 reads from 8 samples (4 eDNA and 4 iDNA), resulting in 167,330 contigs. We predicted 219,893 open reading frames (ORFs), of which approximately 66% could be annotated using Cluster of Orthologous Genes (COGs) and around 49% had Kyoto Encyclopedia of Genes and Genomes (KEGG) annotations. COG annotations were used to determine the broader functions represented in the metagenome and metatranscriptome, while KEGG annotations were used to elucidate the metabolic networks in the DWDS microbiome. The mapping percentage for iDNA sequences was about 97%, compared to approximately 96% for eDNA samples.

**Text S3:**

***Regulating cell membrane structure.*** An essential aspect of survival in the DWDS is the regulation and maintenance of cell membrane properties. We found that the synthesis of membrane components like phospholipids was preferred from triglycerides rather than from acyl-ACP produced through fatty acid biosynthesis (Figure S13). The phospholipid transporter, which was enriched in iRNA, plays a crucial role in maintaining membrane structure and thus is vital for cell survival (Figure 5A)[1]. Other membrane components, such as peptidoglycans, are synthesized by cross-linking uridine di-phosphate N-acetylglucosamine (UDP-GlcNAc) and UDP-N-acetylmuramic acid (UDP-MurNAc). Transcripts for these compounds were highly enriched in the iRNA (Figures 5B, S14, and S15). Fructose primarily serves as a precursor for the synthesis of GlcNAc and MurNAc, while glucose and mannose are used to produce other sugar moieties. These sugars are converted into 6-phosphate sugars (e.g., glucose to glucose 6-phosphate), which are then transformed into 1- phosphate sugars. In the oligotrophic environment of the DWDS, organisms reduced the flux toward the production of 1-phosphate sugars (with transcripts enriched in eRNA). Instead, the 6-phosphate sugars were isomerized to fructose-6-phosphate, which was then used to produce UDP-GlcNAc and UDP-MurNAc. UDP-GlcNAc further contributes to the formation of capsular and O-antigens building blocks such as UDP-N-acetyl-D-galactosaminuronic acid and UDP-N-acetyl-D-glucosaminuronate[2,3]. Glucose-6-phosphate, which converts to glucose 1-phosphate, is preferentially used for the synthesis of polyketide dTDP-L-rhamnose or dTDP-4-amino-4,6-dideoxy-D-galactose (dTDP-Fuc4N) instead of being converted into amylose for storage. Both dTDP-L-rhamnose and dTDP-Fuc4N are used for synthesis of surface polysaccharides.

**Text S4:**

***Carbon metabolism and energy generation.*** Efficient energy generation is crucial in the DWDS. Genes involved in carbon and amino acid metabolism pathways (Figures S16, and S17), such as glycolysis, pyruvate metabolism, and the TCA cycle, were enriched in the iRNA (Figure 4A). A closer examination of these pathways reveals that only the genes responsible for the oxidation of pyruvate to acetyl-CoA were upregulated for the iRNA fraction (Figure 5C). Acetyl-CoA enters the TCA cycle to produce ATP and other energy-rich molecules (NADH, $FADH_2$, and GTP), which are converted into ATP through oxidative phosphorylation (with multiple genes encoding different subunits enriched in iRNA, Figure S15). Additionally, lactate and methylglyoxal (a toxic compound) can also be converted to pyruvate for further energy generation. The conversion of S-malate into Malyl-CoA was less favored (enriched in eRNA), highlighting the importance of retaining S-malate in the TCA cycle. The accumulation of phosphoenolpyruvate (PEP) was promoted and subsequently utilized for the synthesis of chorismate via the shikimate pathway (Figure 5C). Chorismate is preferentially utilized for the synthesis of ubiquinones (which facilitates energy generation[4]) rather than being converted into tryptophan, tyrosine, or phenylalanine. Furthermore, the synthesis of L-serine was enriched in eRNA, indicating an effort to maintain the flow of glycerate 3-phosphate within glycolysis for the formation of pyruvate and subsequent energy generation.

**References**


1. Shrivastava, R. & Chng, S.-S. Lipid trafficking across the Gram-negative cell envelope. *Journal of Biological Chemistry* **294**, 14175–14184 (2019).
2. Crépin, S. *et al.* The UDP-GalNAcA biosynthesis genes gna-gne2 are required to maintain cell envelope integrity and in vivo fitness in multi-drug resistant Acinetobacter baumannii. *Molecular Microbiology* **113**, 153–172 (2020).
3. Zhao, X. *et al.* WbpO, a UDP-galactosamine Dehydrogenase from Pseudomonas aeruginosa Serotype O6 *. *Journal of Biological Chemistry* **275**, 33252–33259 (2000).
4. Pierrel, F., Burgardt, A., Lee, J.-H., Pelosi, L. & Wendisch, V. F. Recent advances in the metabolic pathways and microbial production of coenzyme Q. *World Journal of Microbiology and Biotechnology* **38**, 58 (2022).

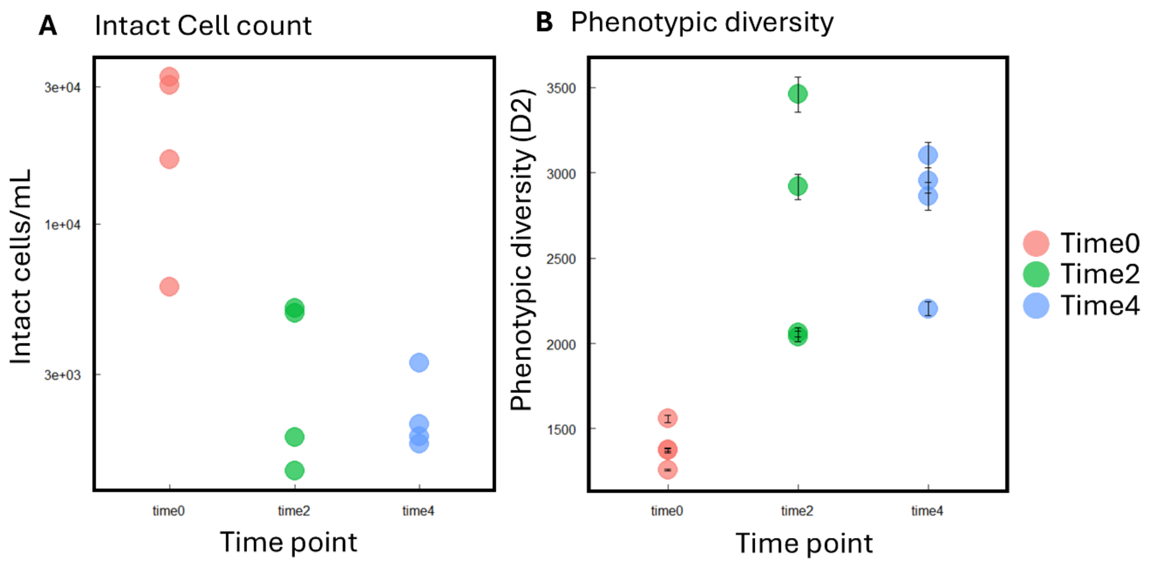


**Figure S1.** A) Intact cell counts at start (time0), middle (time2), and end (time4) points of the sampling process. B) Phenotypic diversity represented by 2nd order Hill number (D2) shows increased diversity of drinking water with continued sampling.

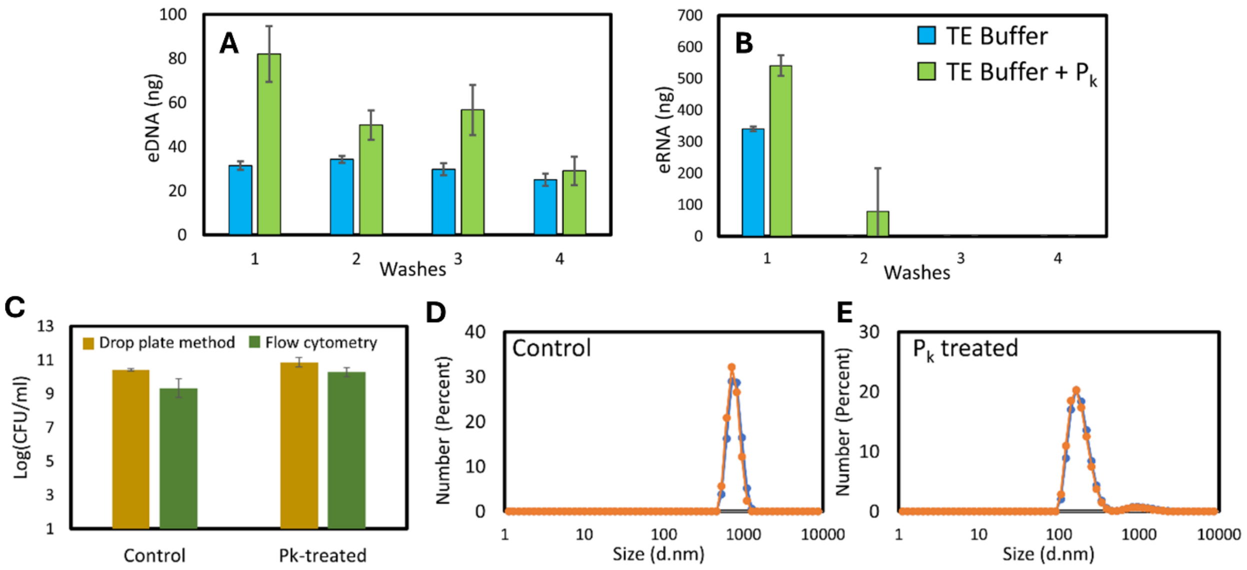


**Figure S2.** Optimization of eNAC extraction from DWDS. A) eDNA and B) eRNA recovery from filter samples with and without proteinase K treatment using TE buffer as extraction buffer (n = 4). C) Proteinase K treatment has no effect on the viability and permeability of the cells. Viability measured using drop plate method and Flow cytometry using Live dead staining (n=3). Particle Size analysis shows shift of peak to smaller size particles. D) Particle size in control samples. E) Particle size after $P_k$ treatment. Different lines indicate replicate measurement.

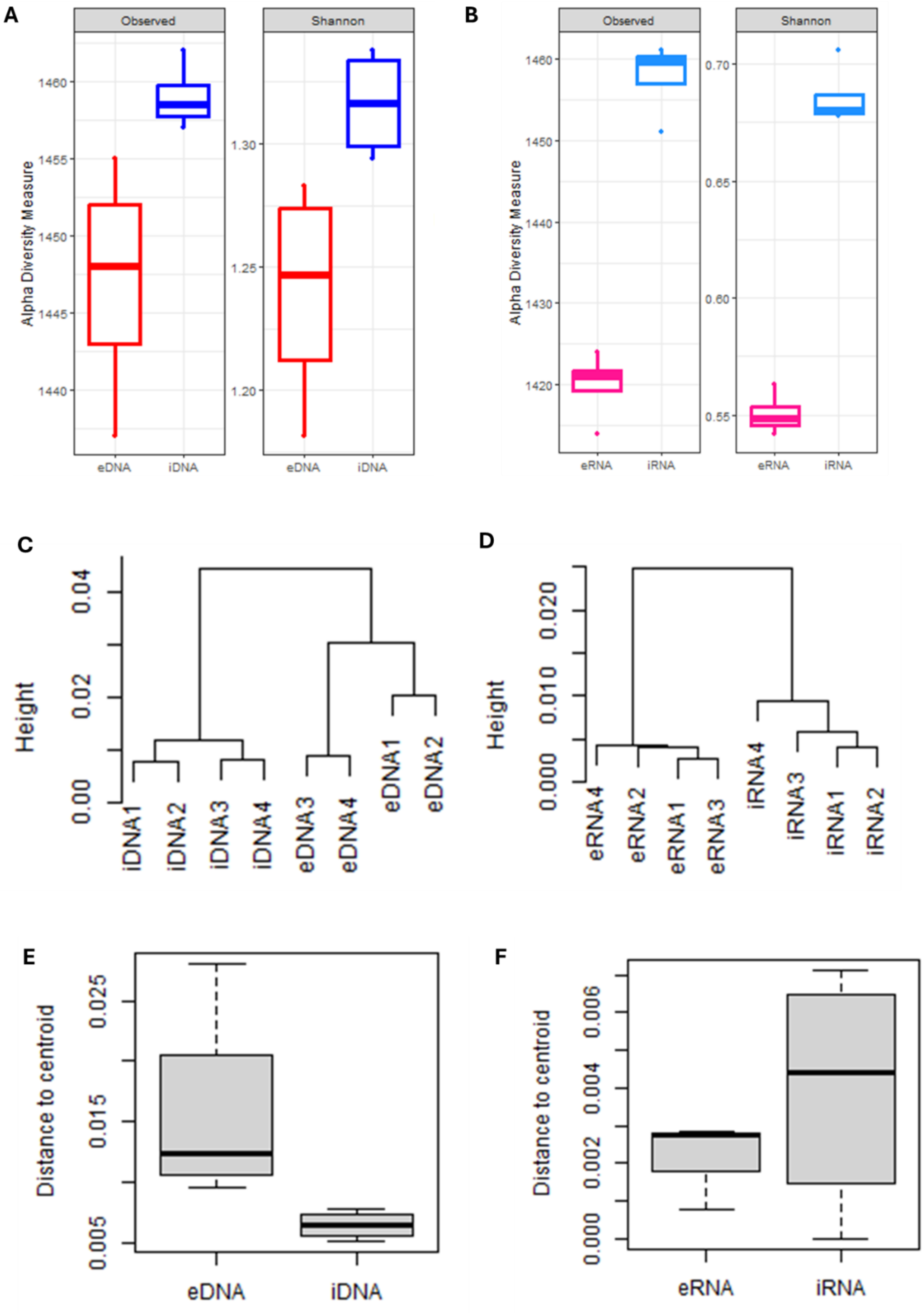


**Figure S3.** The α-diversity (local microbial diversity at the sampling location) for microbial communities detected by (A) DNA and (B) RNA sequencing. Hierarchical clustering of the samples for DNA (C) and RNA (D) based on the average linkage clustering method. The branch length describes the difference between the samples. Difference in dispersion between E) eDNA and iDNA, and F) eRNA and iRNA samples.

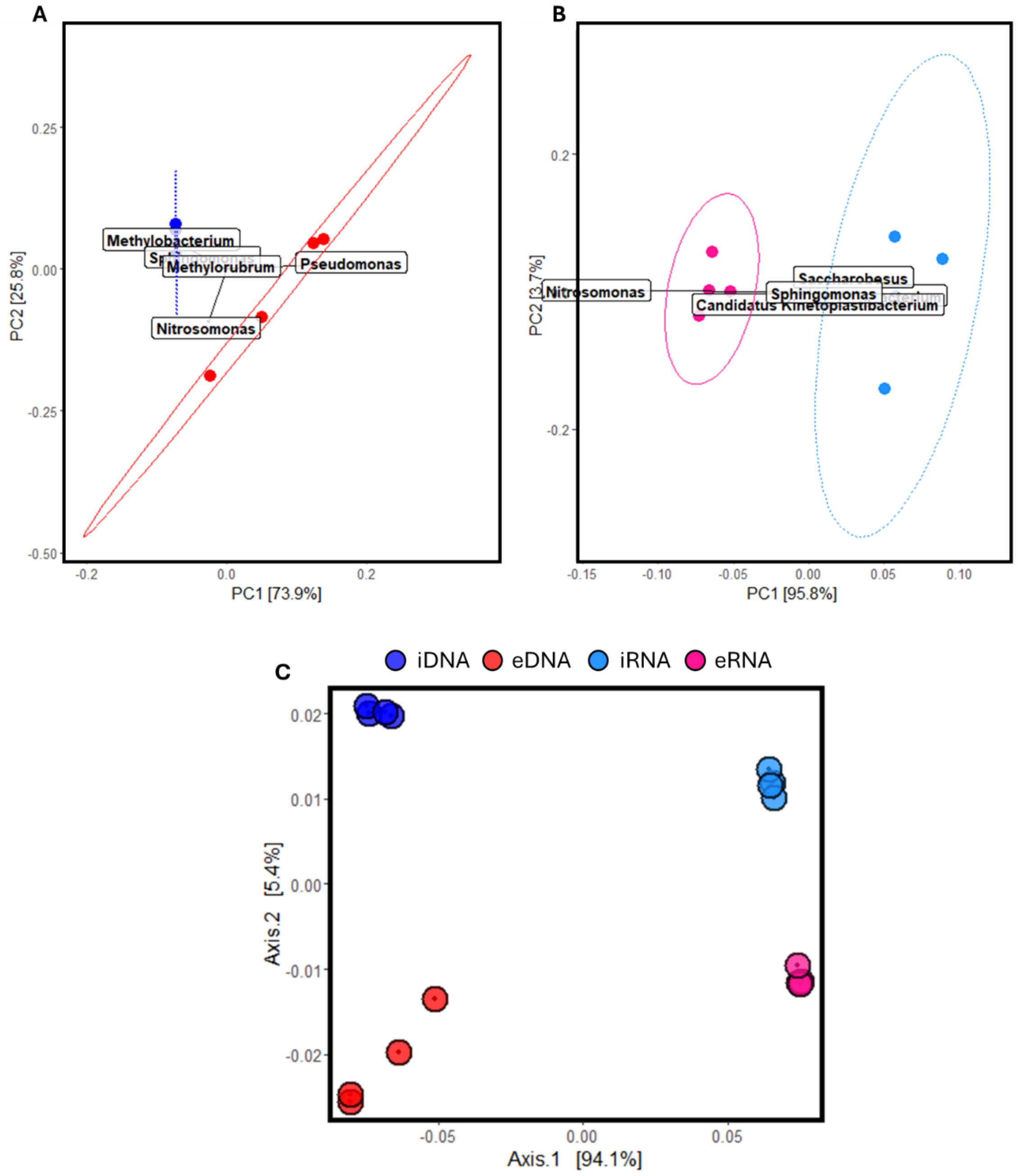


**Figure S4.** PCoA plot showing the difference between communities detected in the nucleic acid fractions based on Bray-Curtis dissimilarity.

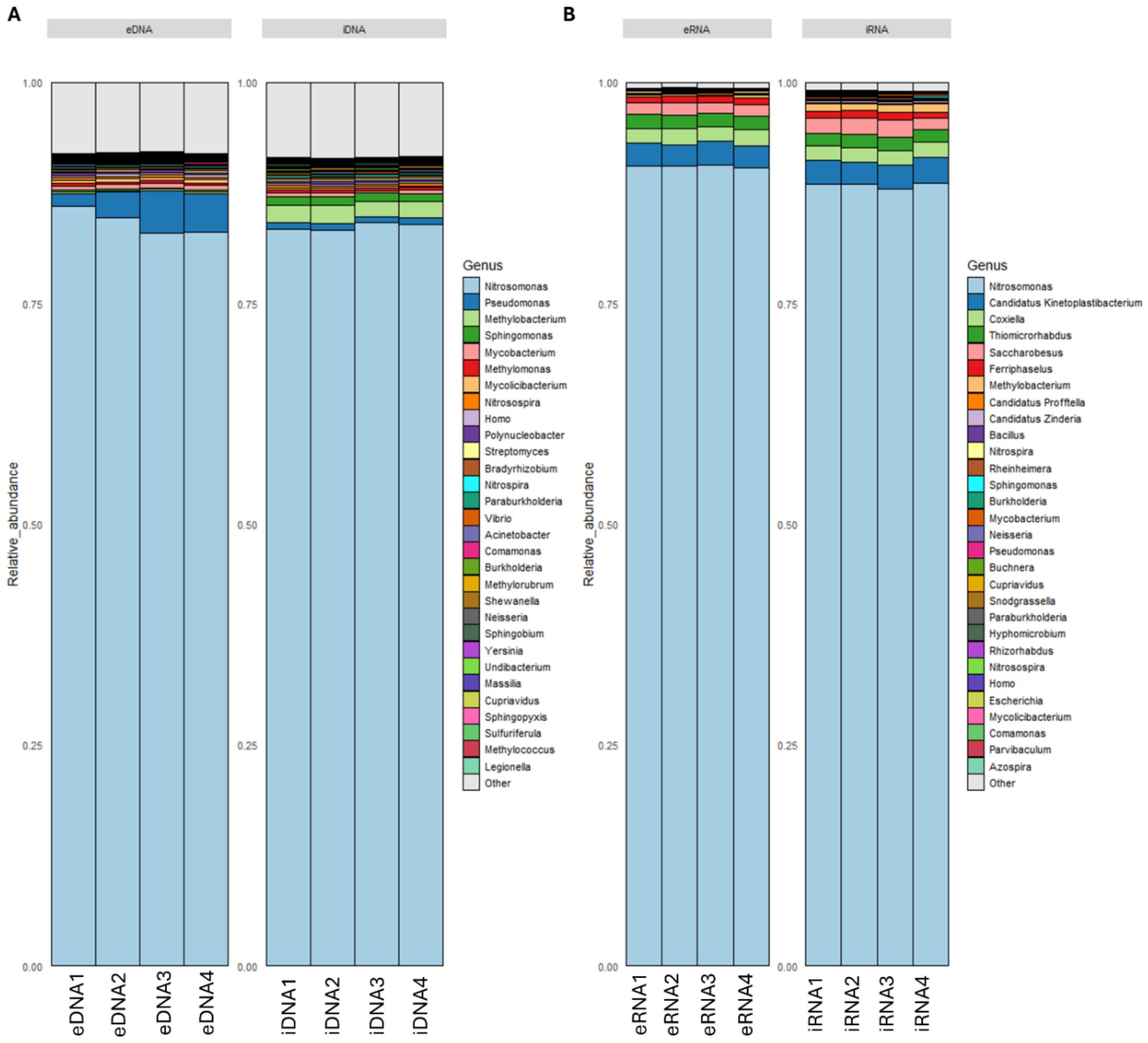


**Figure S5.** Community composition at genus level for A) DNA, and B) RNA. Relative abundance from top 30 genera were selected as they reflect major genera detected.

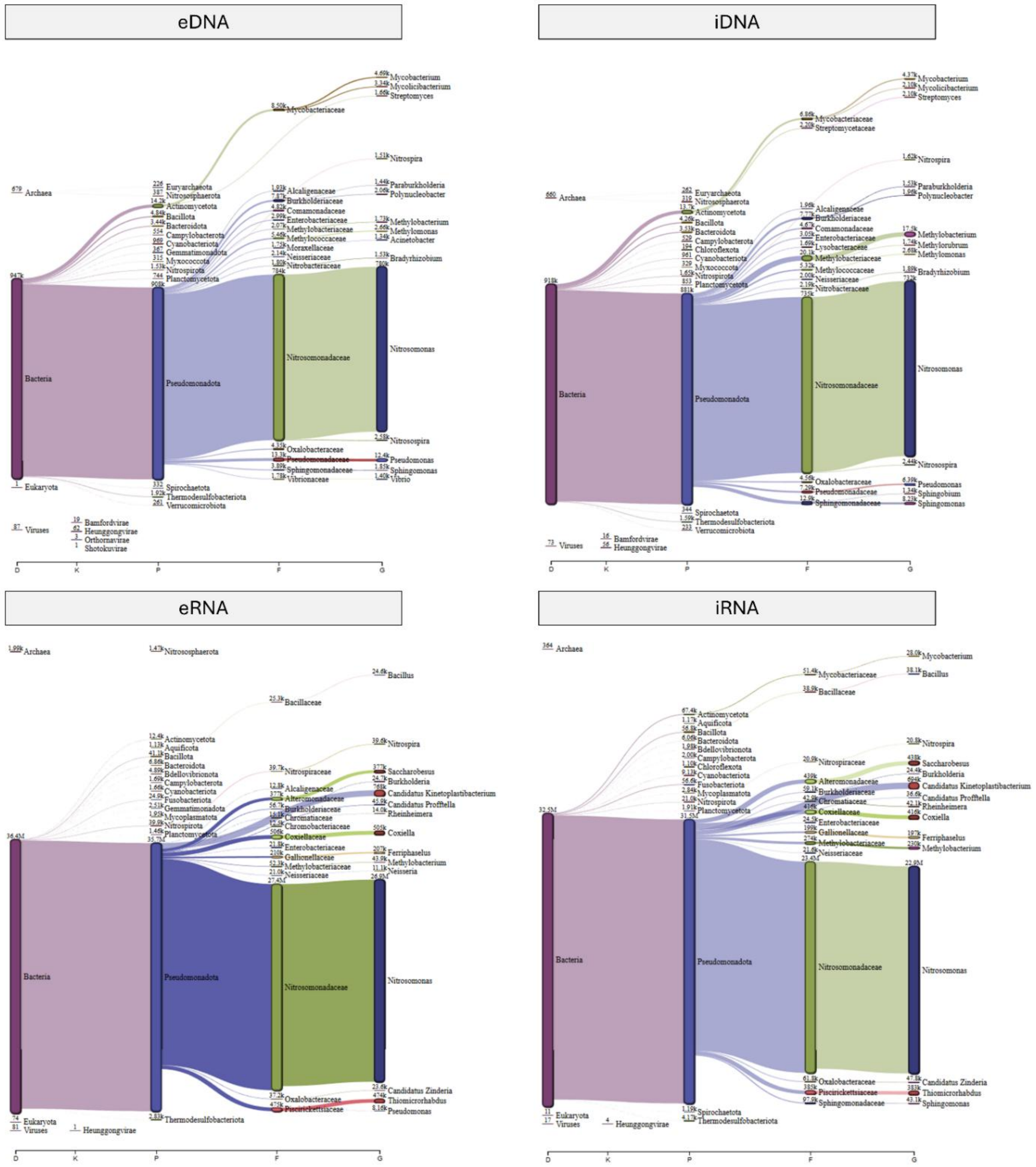


**Figure S6.** Snakey visualization of assigned reads in eDNA, iDNA, eRNA, and iRNA. The representative images are from sample 1 of each nucleic acid fraction.

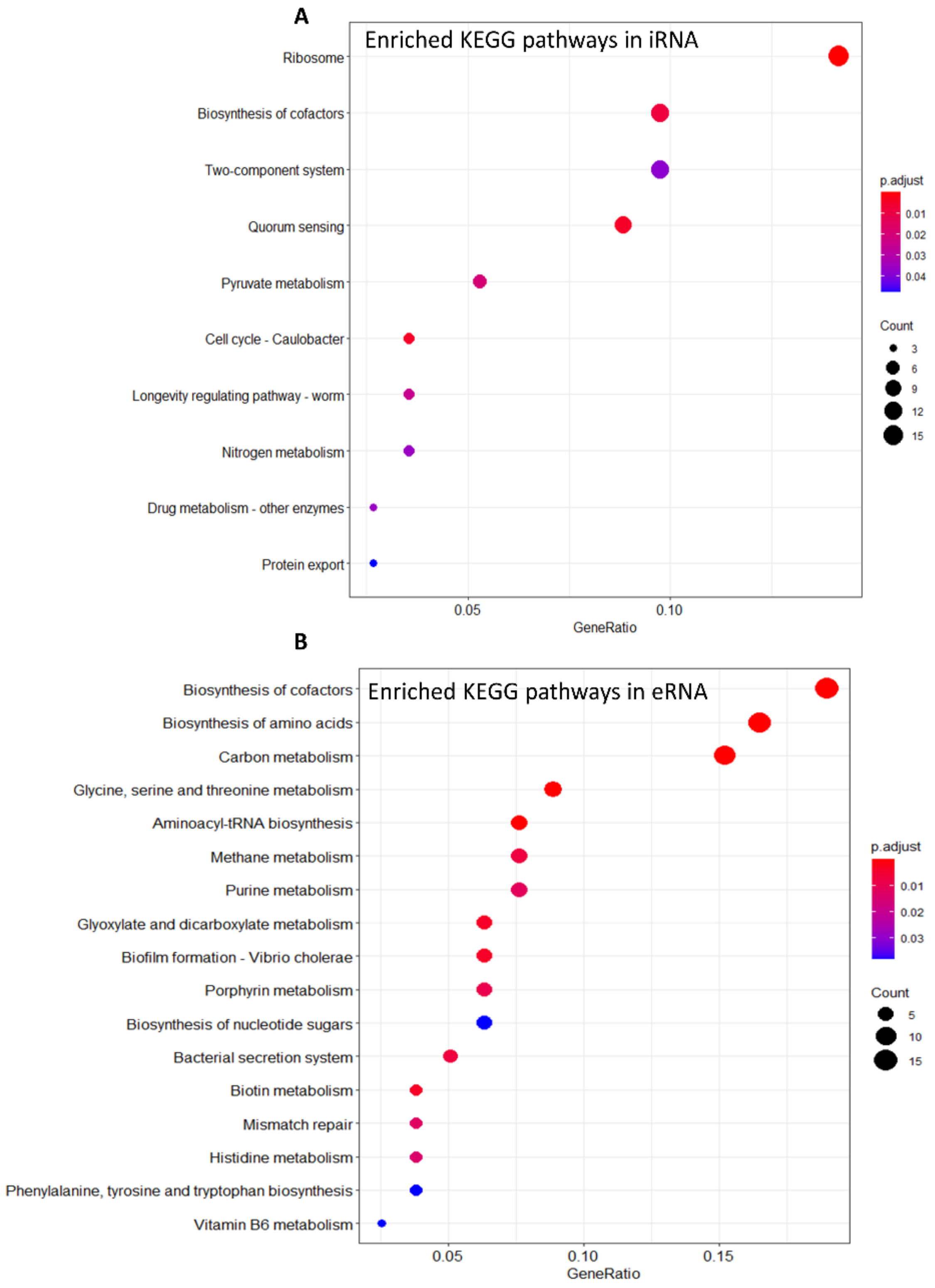


**Figure S7.** Pathways represented by significantly enriched genes in A) iRNA and B) eRNA

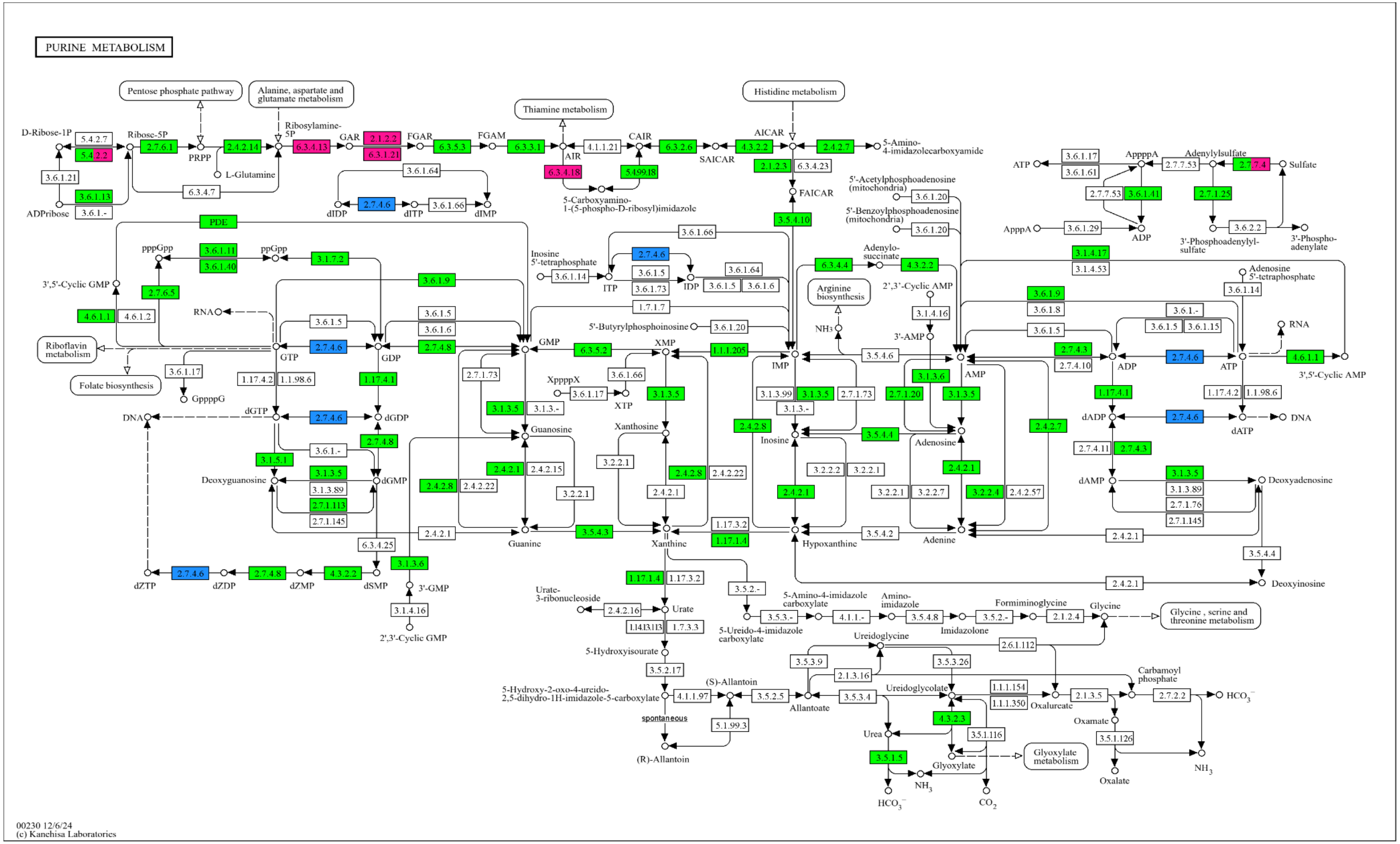


**Figure S8.** KEGG pathway for purine metabolism. Green represents enzymes/pathways not significantly enriched in iRNA and eRNA, light blue represents enzymes/pathways significantly enriched in iRNA while pink represents enzymes/pathways significantly enriched in eRNA (with p-adjusted < 0.01 and $|\log_2(\text{fold-change})| > 0.58$)

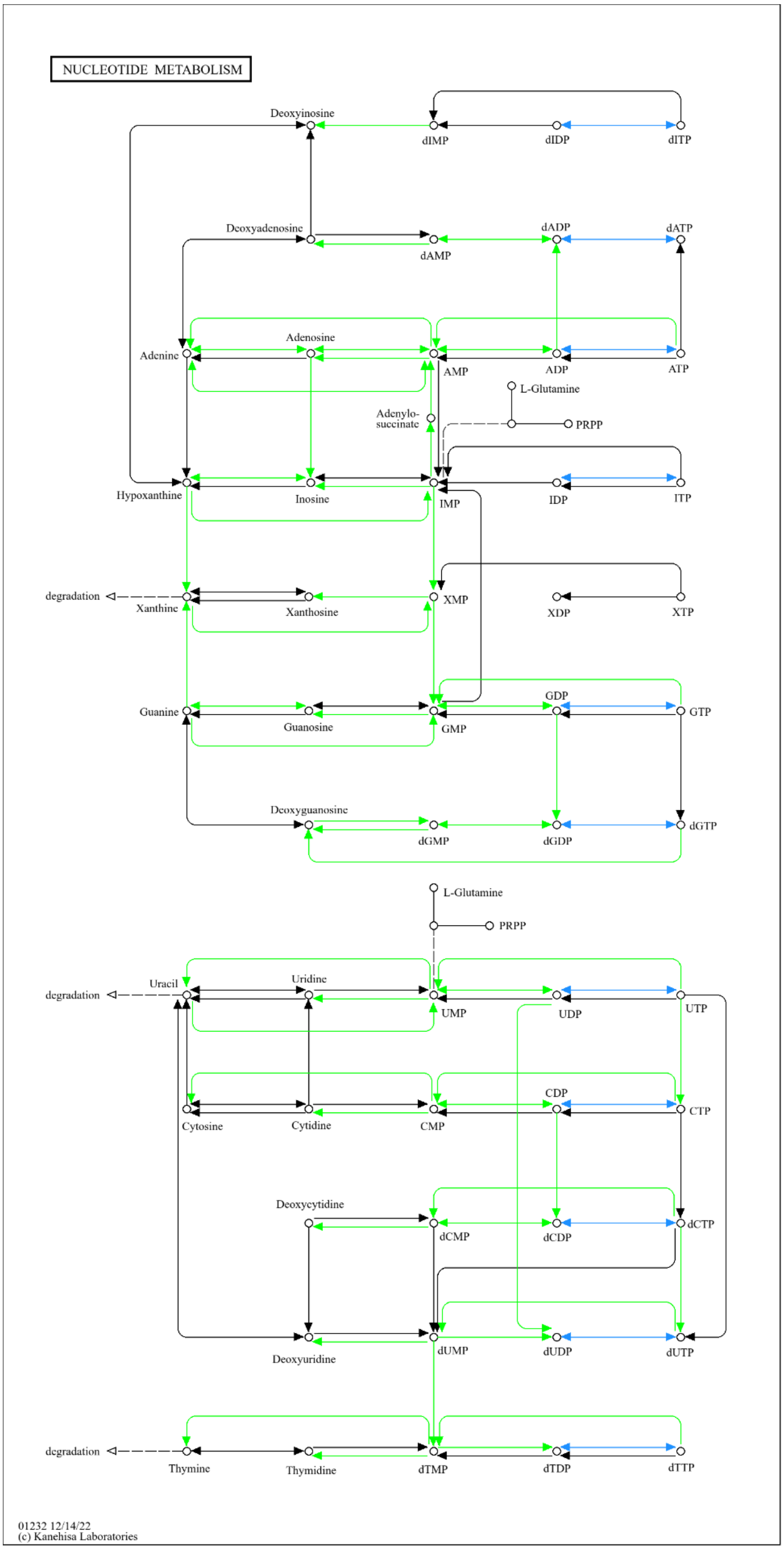


**Figure S9.** KEGG pathway for nucleotide metabolism. Green represents enzymes/pathways not significantly enriched in iRNA and eRNA, light blue represents enzymes/pathways significantly enriched in iRNA while pink represents enzymes/pathways significantly enriched in eRNA (with p-adjusted < 0.01 and $|\log_2(\text{fold-change})| > 0.58$)

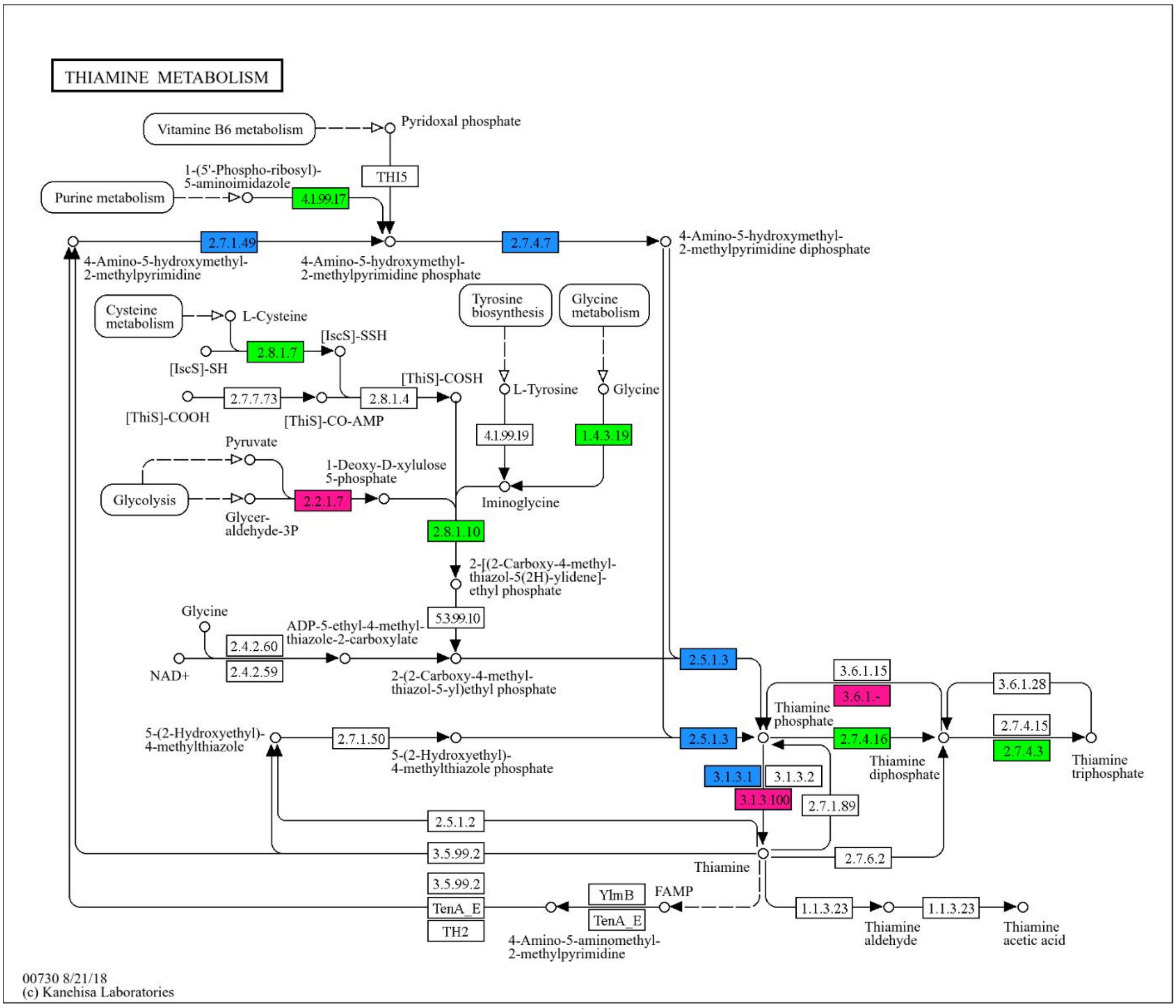


**Figure S10.** KEGG pathway for thiamine metabolism. Green represents enzymes/pathways not significantly enriched in iRNA and eRNA, light blue represents enzymes/pathways significantly enriched in iRNA while pink represents enzymes/pathways significantly enriched in eRNA (with p-adjusted $< 0.01$ and $|\log_2(\text{fold-change})| > 0.58$)

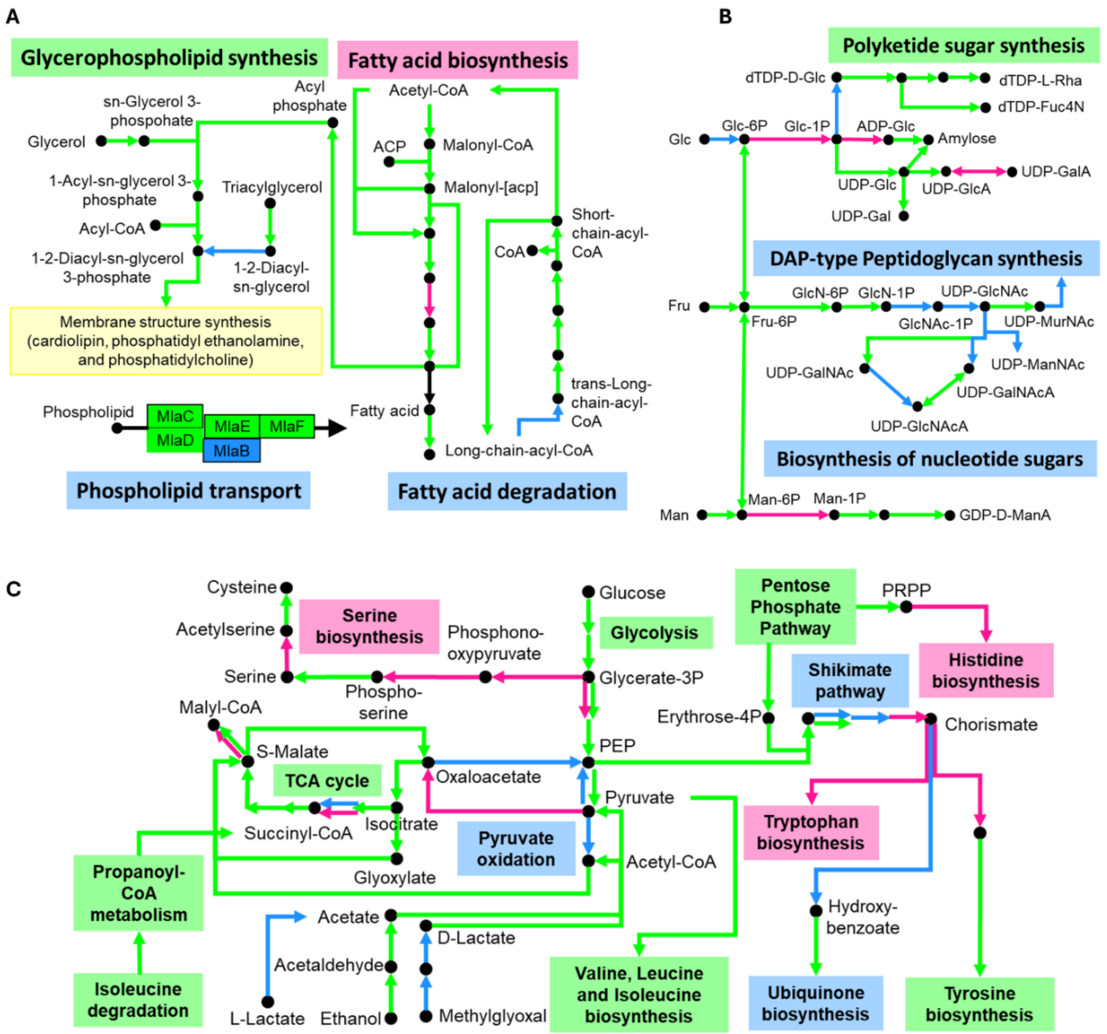


**Figure S11. Pathway flux within different pathways in the DWDS. A)** Fatty acid degradation is preferred over fatty acid synthesis while membrane components are derived from triacylglycerols. Phospholipid transporters help maintain and modify cell membrane structure according to environmental conditions. **B)** Sugars like glucose (Glc), fructose (Fru) and mannose (Man) are preferentially utilized for synthesis of Uridine diphosphate-N-acetylglucosamine (UDP-GlcNAc) and UDP-N-acetylmuramic acid (UDP-MurNAc) which are used in peptidoglycan biosynthesis **C)** Major energy generation is through glycolysis (glucose converted to 2*pyruvate) and tricarboxylic acid (TCA) cycle. Ethanol and Methylglyoxal are also converted to pyruvate which is preferentially utilized by conversion into acetyl CoA which is fed into the TCA cycle. Phosphoenolpyruvate (PEP) is channeled into the shikimate pathway for production of chorismite which is the preferentially utilized for synthesis of ubiquinones over amino acid biosynthesis. Green represents enzymes/pathways not significantly enriched in iRNA and eRNA, light blue represents enzymes/pathways significantly enriched in iRNA while pink represents enzymes/pathways significantly enriched in eRNA (with p-adjusted $< 0.01$ and $|\log_2(\text{fold-change})| > 0.58$). **Black arrows** represent enzymes/pathways **not detected** in the current data. ACP: Acyl-carrier protein; dTDP-D-Glc: 2'Deoxy-thymidine-5'-diphospho-alpha-D-glucose; dTDP-L-Rha: dTDP-L-Rhamnose; dTDP-Fuc4N: dTDP-4-amino-4,6-dideoxy-D-galactose; UDP-GlcA: UDP-glucuronate; UDP-GalA: UDP-galacturonate; UDP-GlcNAcA: UDP-N-acetyl-D-glucosaminuronate; UDP-GalNAcA: D-galactosaminuronic acid. Multiple lines indicate different genes for the same reaction.

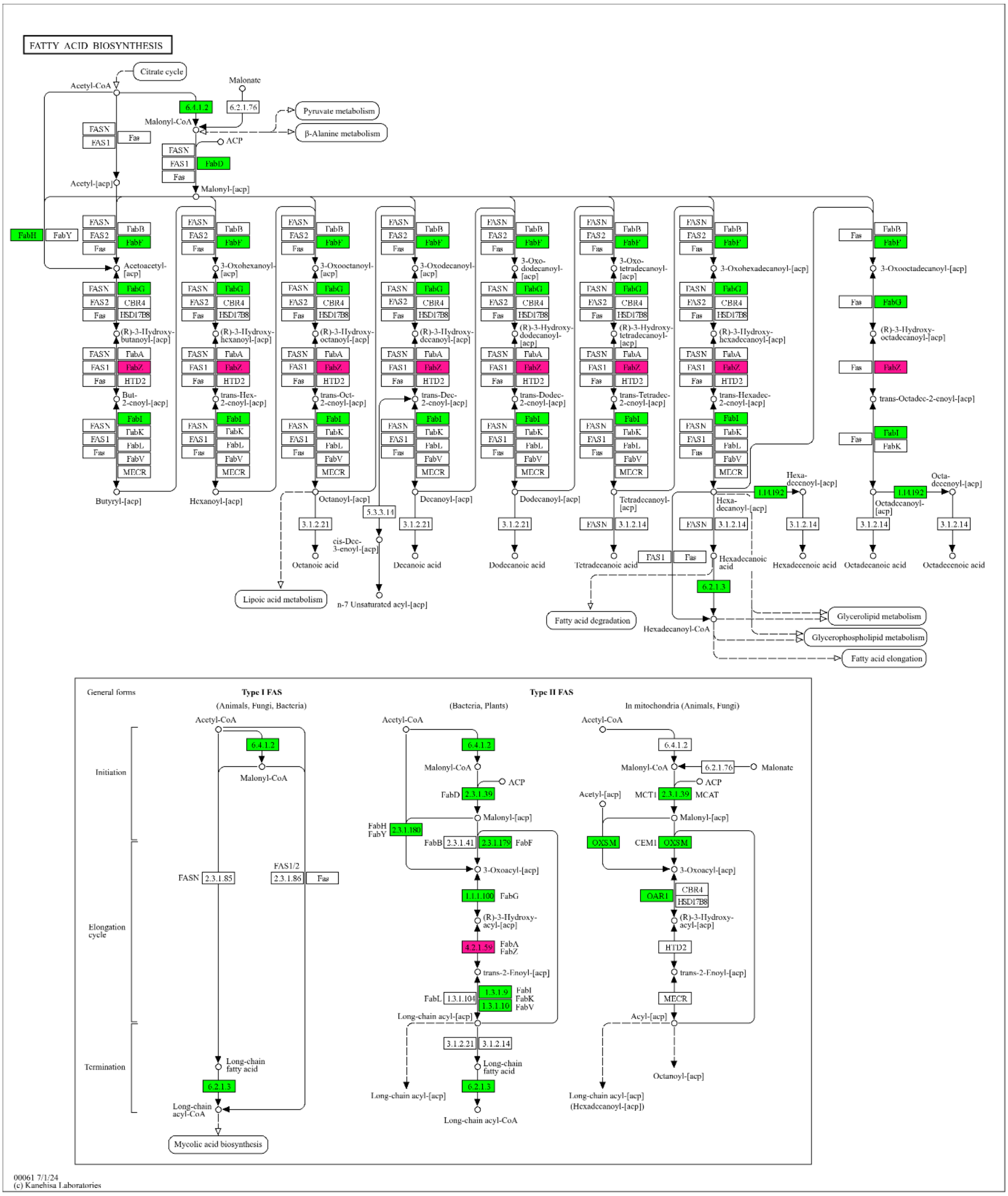


**Figure S12.** KEGG pathway for fatty acid biosynthesis. Green represents enzymes/pathways not significantly enriched in iRNA and eRNA, light blue represents enzymes/pathways significantly enriched in iRNA while pink represents enzymes/pathways significantly enriched in eRNA (with p-adjusted $< 0.01$ and $|\log_2(\text{fold-change})| > 0.58$)

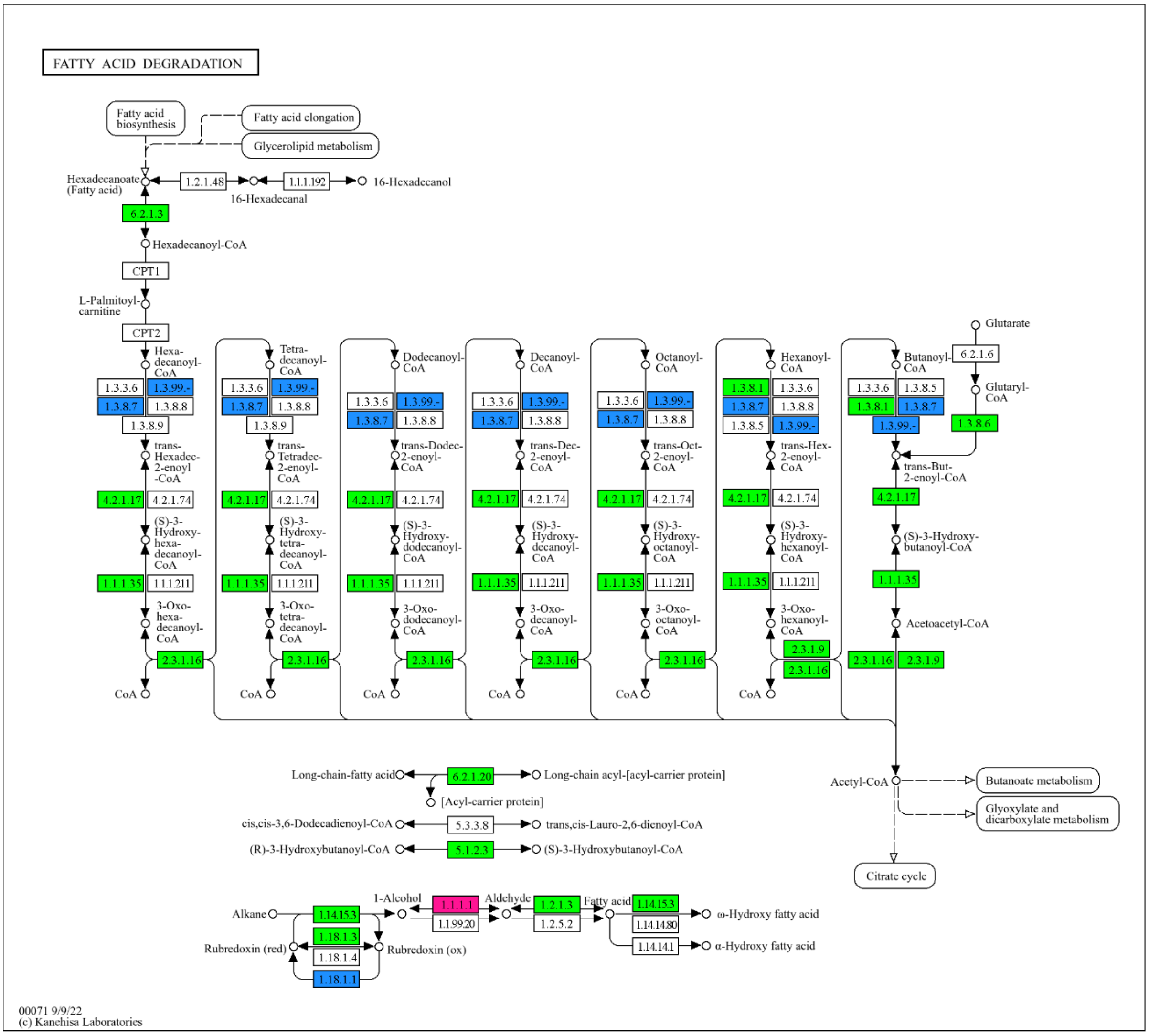


**Figure S13.** KEGG pathway for fatty acid degradation. Green represents enzymes/pathways not significantly enriched in iRNA and eRNA, light blue represents enzymes/pathways significantly enriched in iRNA while pink represents enzymes/pathways significantly enriched in eRNA (with p-adjusted < 0.01 and $|\log_2(\text{fold-change})| > 0.58$)

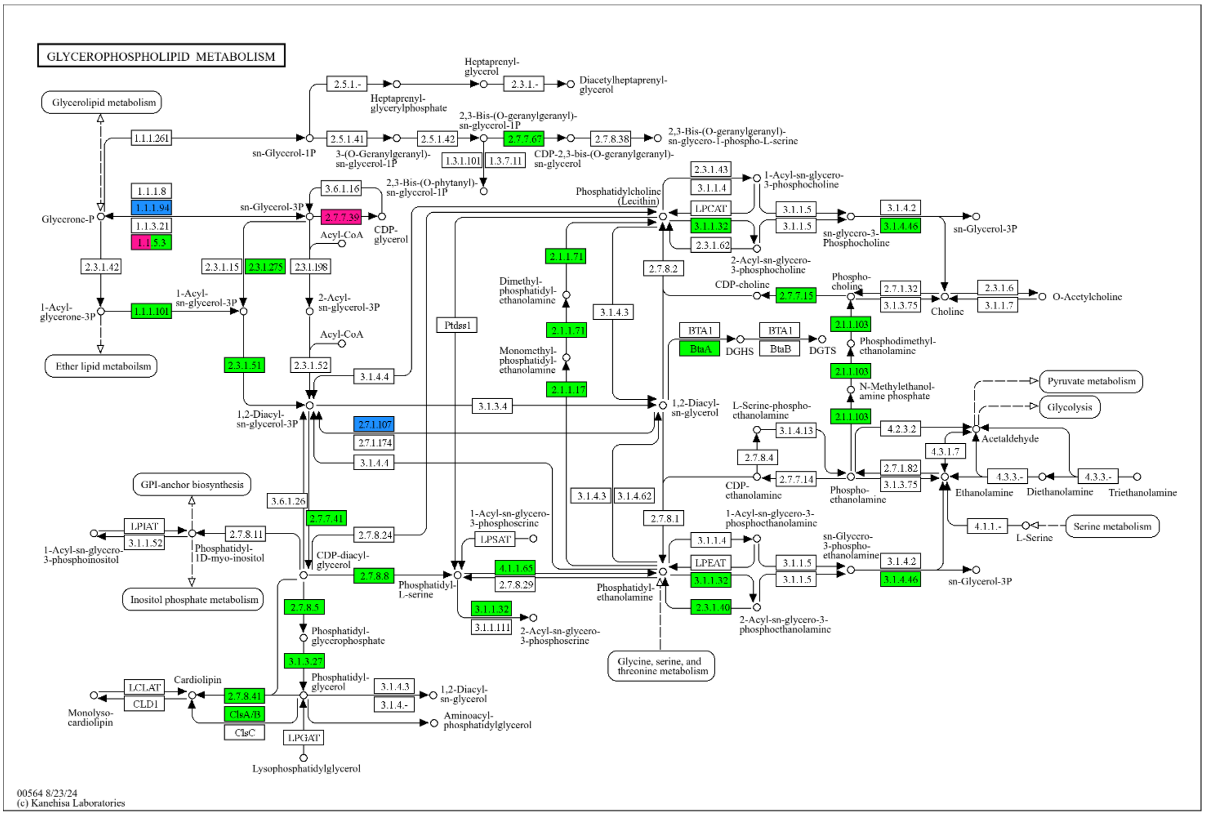


**Figure S14.** KEGG pathway for glycerophospholipid metabolism. Green represents enzymes/pathways not significantly enriched in iRNA and eRNA, light blue represents enzymes/pathways significantly enriched in iRNA while pink represents enzymes/pathways significantly enriched in eRNA (with p-adjusted < 0.01 and $|\log_2(\text{fold-change})| > 0.58$)

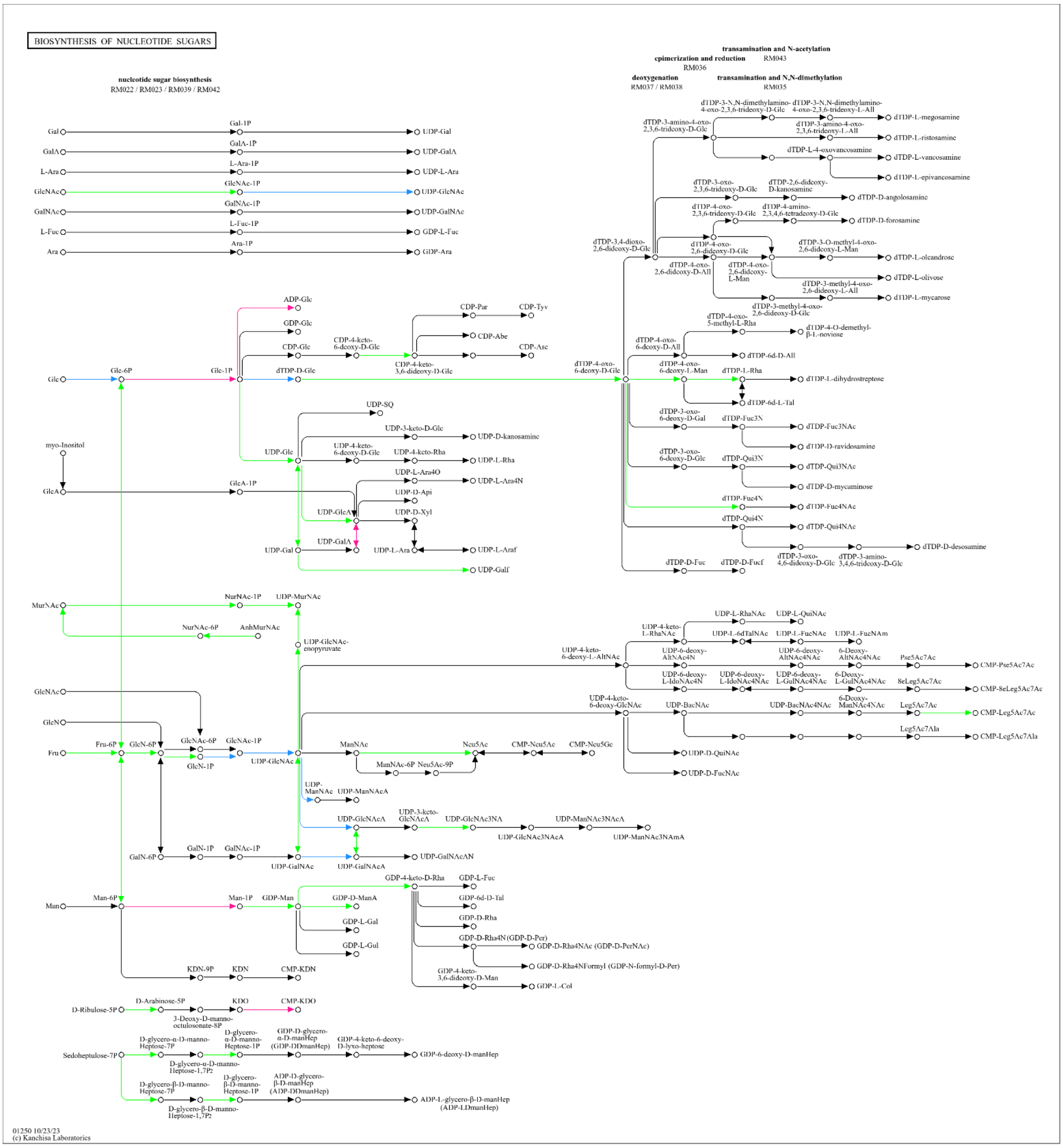


**Figure S15.** KEGG pathway for biosynthesis of nucleotide sugars. Green represents enzymes/pathways not significantly enriched in iRNA and eRNA, light blue represents enzymes/pathways significantly enriched in iRNA while pink represents enzymes/pathways significantly enriched in eRNA (with p-adjusted $< 0.01$ and $|\log_2(\text{fold-change})| > 0.58$)

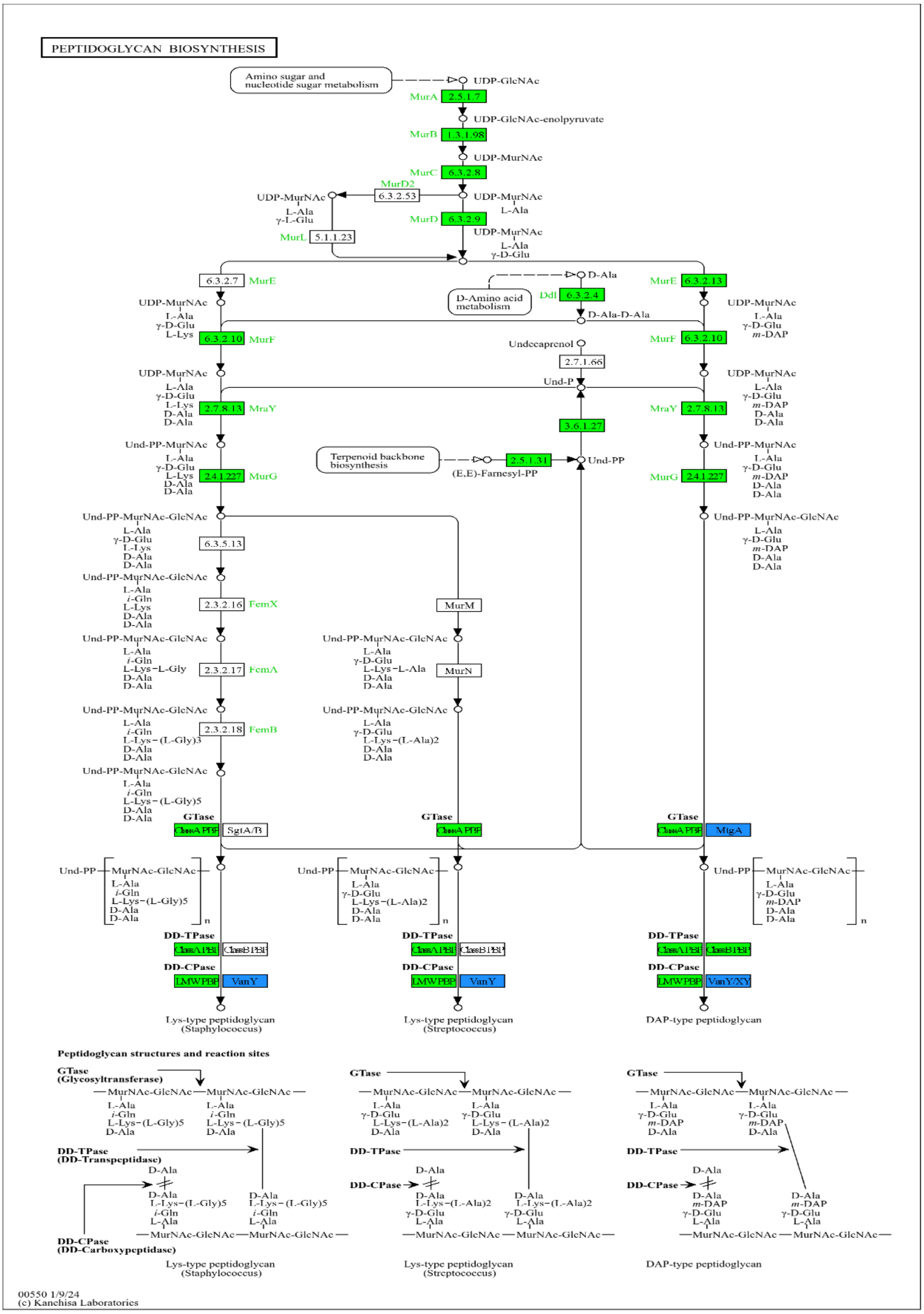


**Figure S16.** KEGG pathway for peptidoglycan biosynthesis. Green represents enzymes/pathways not significantly enriched in iRNA and eRNA, light blue represents enzymes/pathways significantly enriched in iRNA while pink represents enzymes/pathways significantly enriched in eRNA (with p-adjusted $< 0.01$ and $|\log_2(\text{fold-change})| > 0.58$)

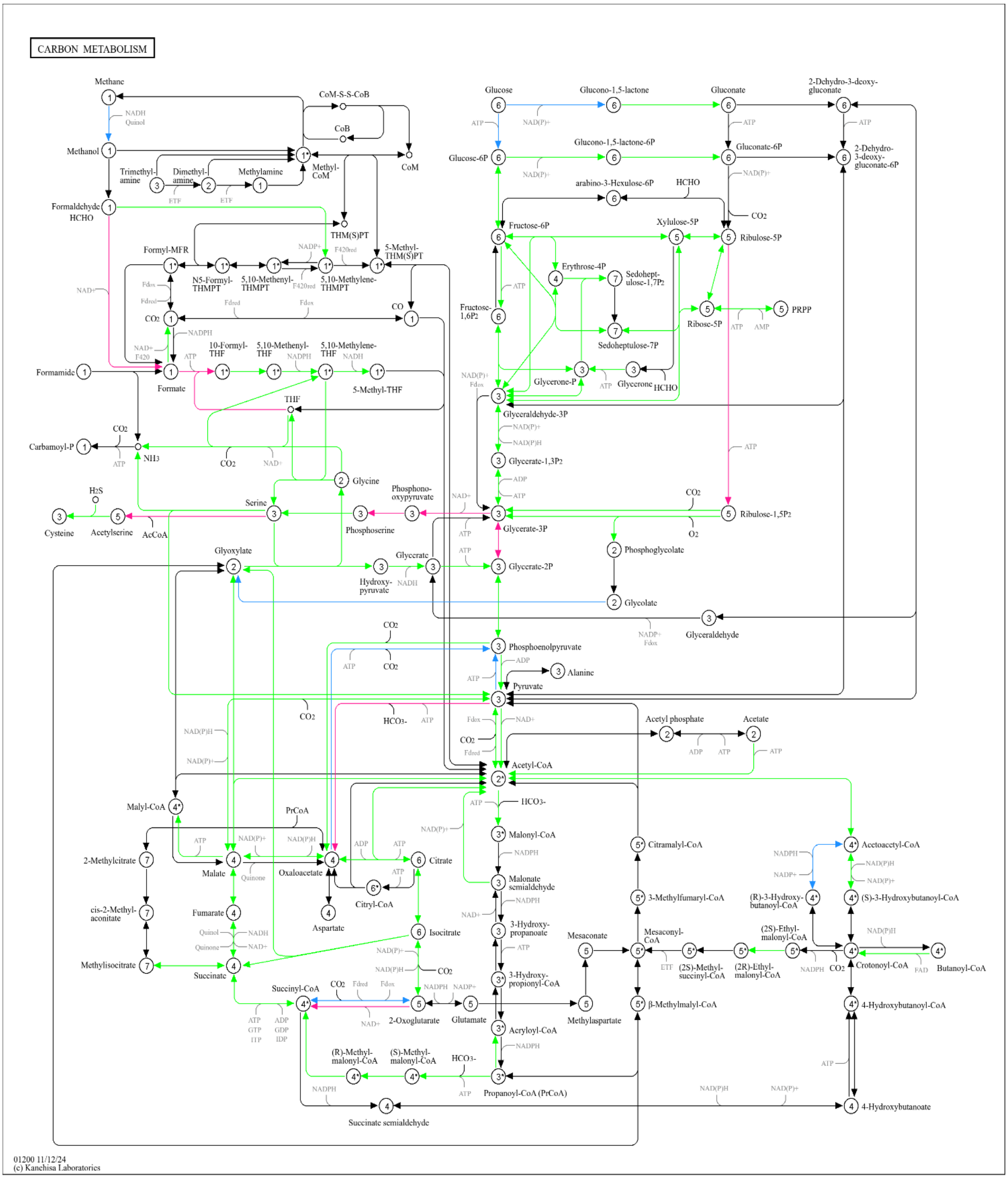

**Figure S17.** KEGG pathway for carbon metabolism. Green represents enzymes/pathways not significantly enriched in iRNA and eRNA, light blue represents enzymes/pathways significantly enriched in iRNA while pink represents enzymes/pathways significantly enriched in eRNA (with p-adjusted < 0.01 and $|\log_2(\text{fold-change})| > 0.58$).

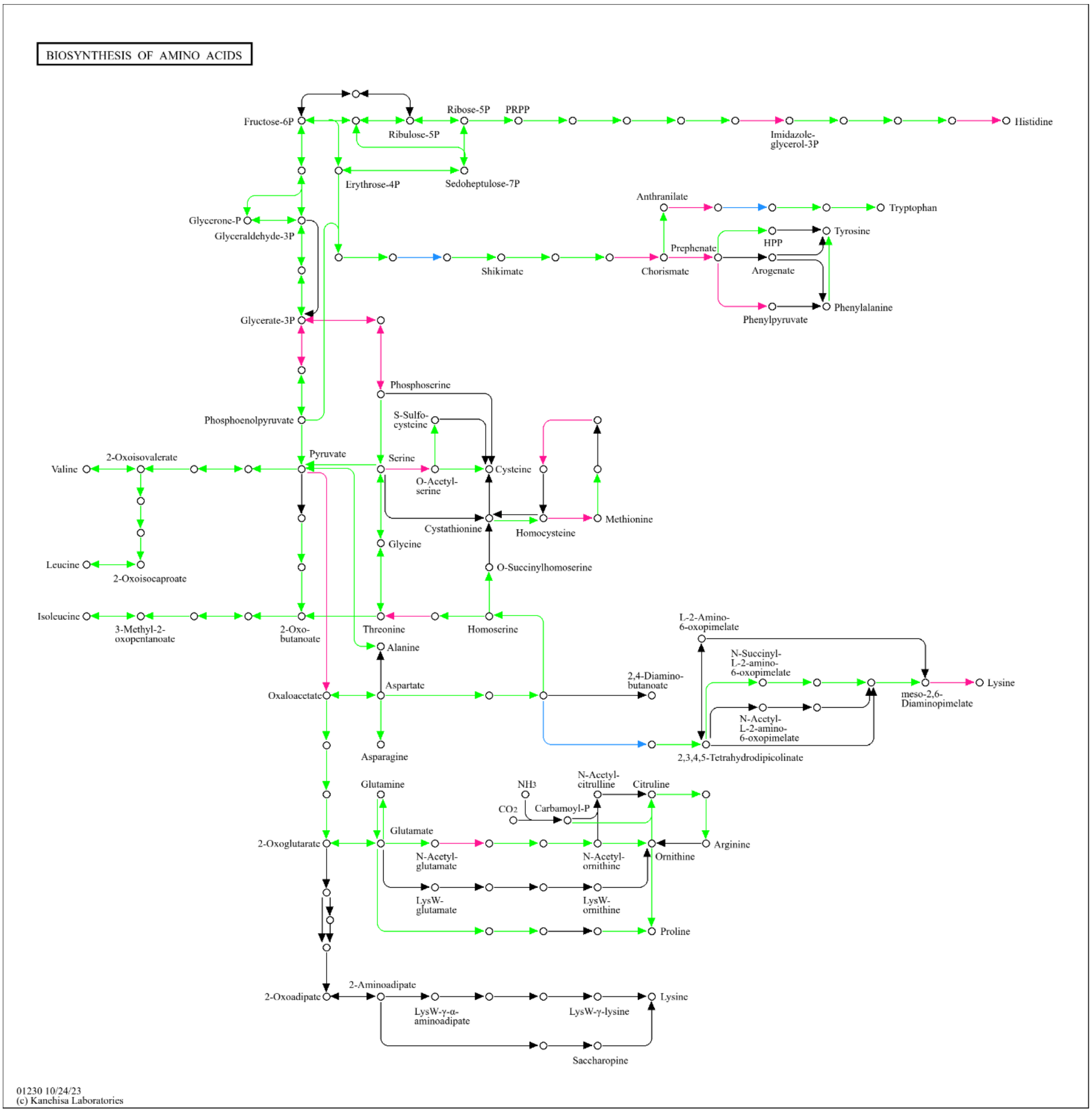


**Figure S18.** KEGG pathway for biosynthesis of amino acids. Green represents enzymes/pathways not significantly enriched in iRNA and eRNA, light blue represents enzymes/pathways significantly enriched in iRNA while pink represents enzymes/pathways significantly enriched in eRNA (with p-adjusted < 0.01 and $|\log_2(\text{fold-change})| > 0.58$).

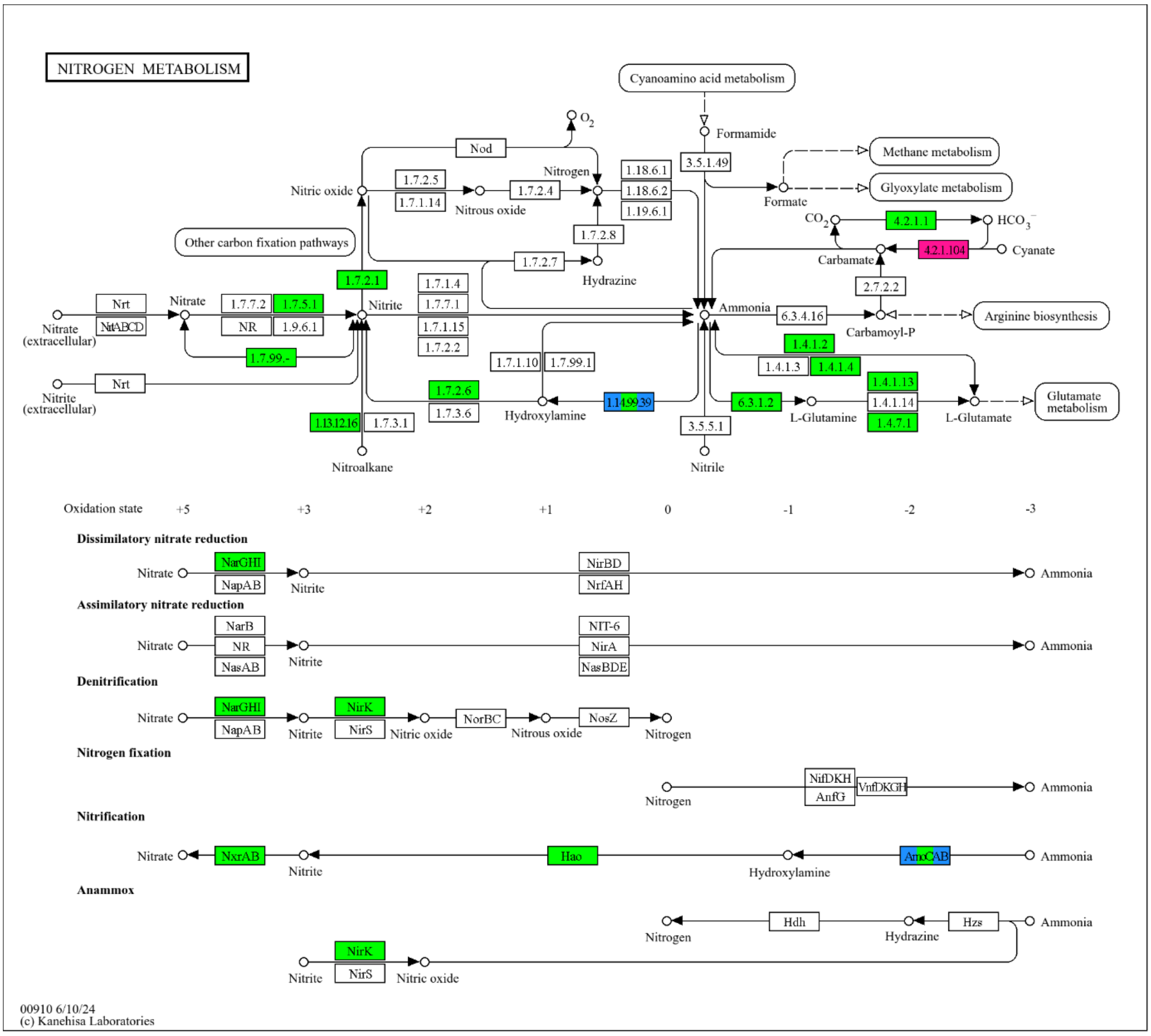


**Figure S19.** KEGG pathway for nitrogen metabolism. Green represents enzymes/pathways not significantly enriched in iRNA and eRNA, light blue represents enzymes/pathways significantly enriched in iRNA while pink represents enzymes/pathways significantly enriched in eRNA (with p-adjusted < 0.01 and $|\log_2(\text{fold-change})| > 0.58$).

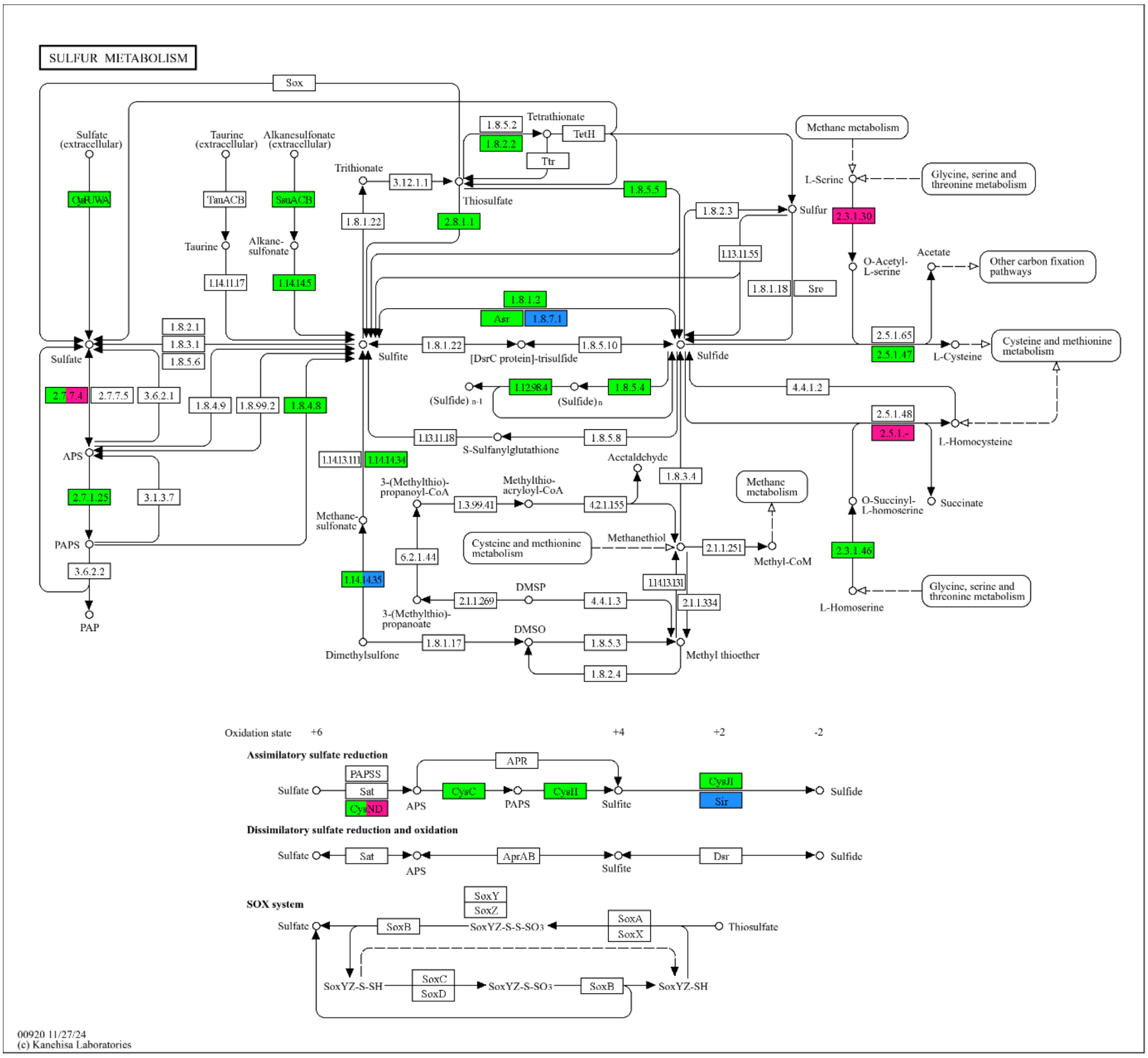


**Figure S20.** KEGG pathway for sulfur metabolism. Green represents enzymes/pathways not significantly enriched in iRNA and eRNA, light blue represents enzymes/pathways significantly enriched in iRNA while pink represents enzymes/pathways significantly enriched in eRNA (with p-adjusted < 0.01 and $|\log_2(\text{fold-change})| > 0.58$).

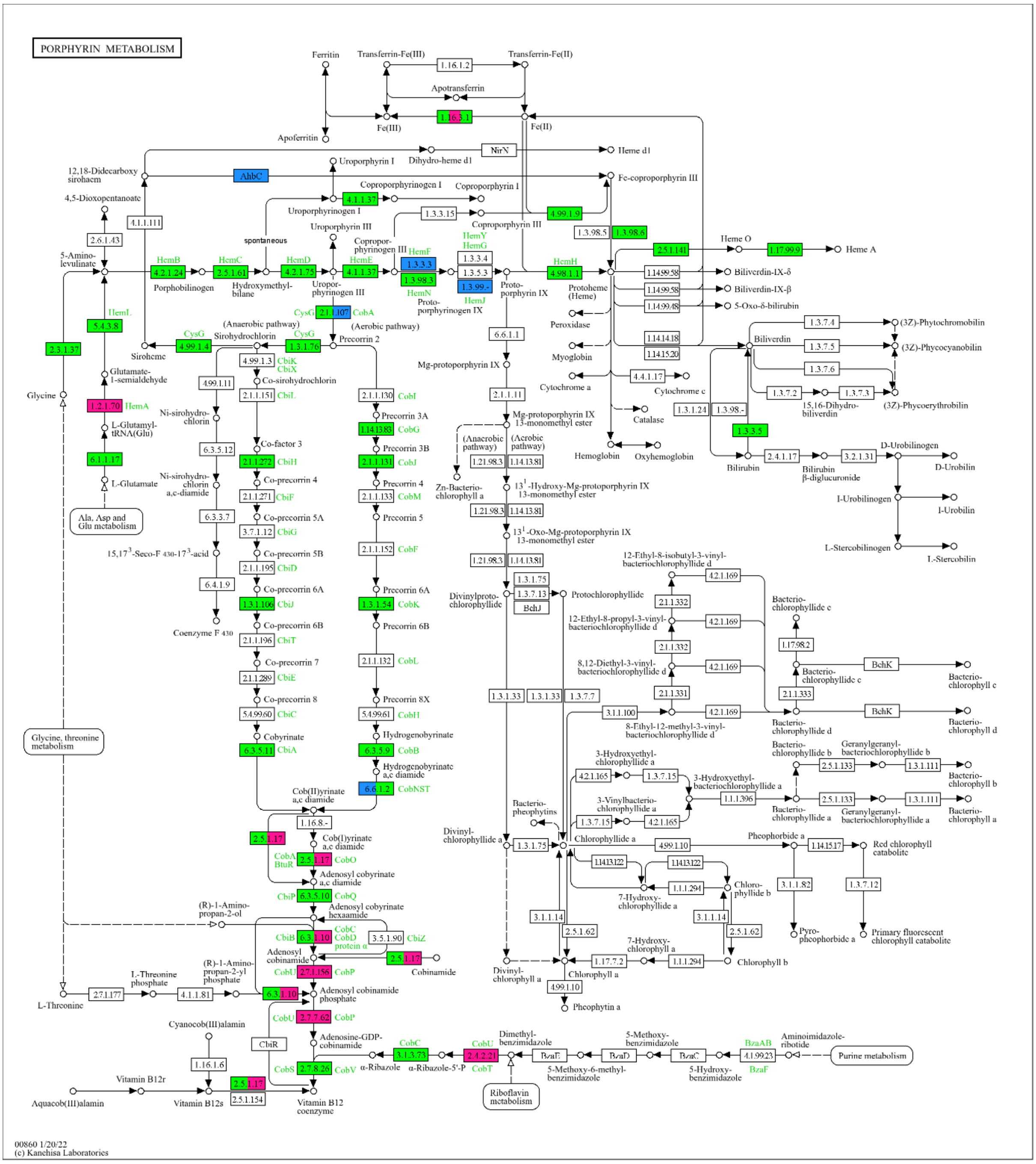

**Figure S21.** KEGG pathway for porphyrin metabolism. Green represents enzymes/pathways not significantly enriched in iRNA and eRNA, light blue represents enzymes/pathways significantly enriched in iRNA while pink represents enzymes/pathways significantly enriched in eRNA (with p-adjusted < 0.01 and $|\log_2(\text{fold-change})| > 0.58$).

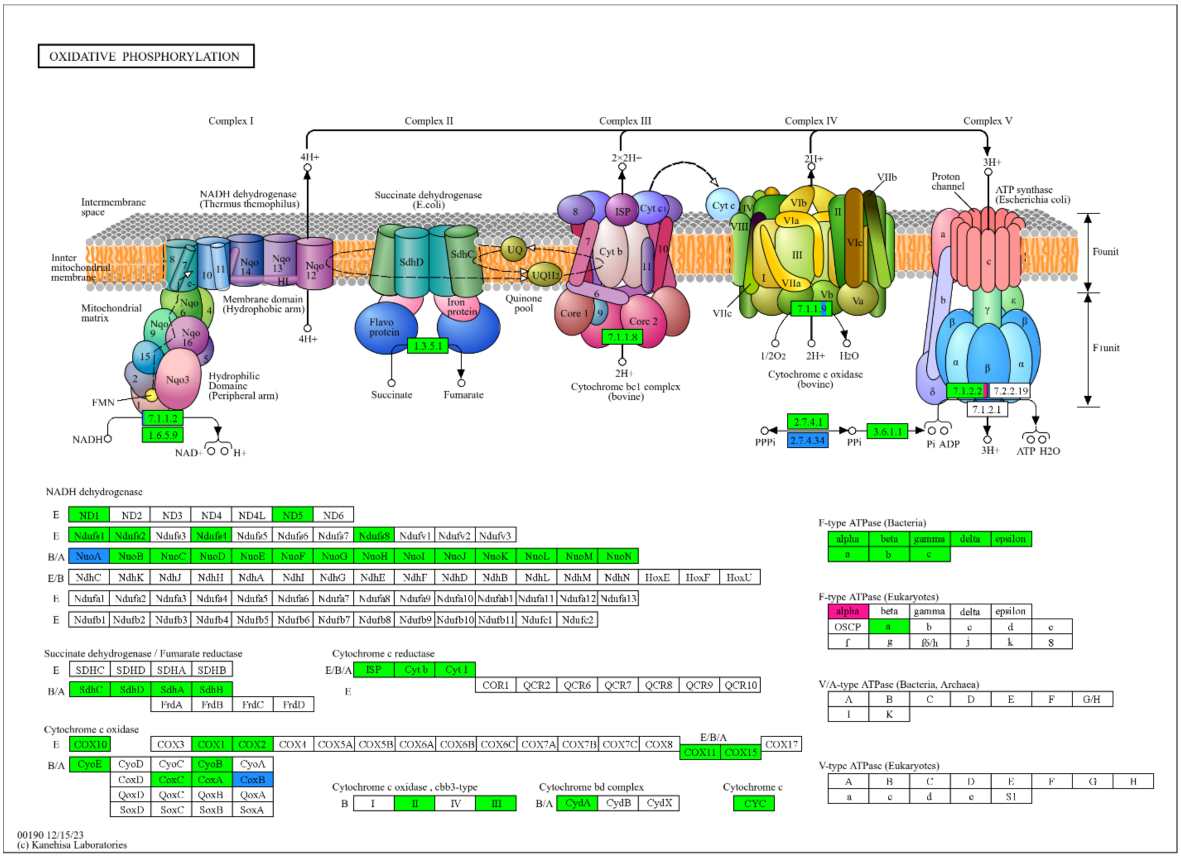


**Figure S22.** KEGG pathway for oxidative phosphorylation. Green represents enzymes/pathways not significantly enriched in iRNA and eRNA, light blue represents enzymes/pathways significantly enriched in iRNA while pink represents enzymes/pathways significantly enriched in eRNA (with p-adjusted < 0.01 and |$\log_2$(fold-change)| > 0.58).

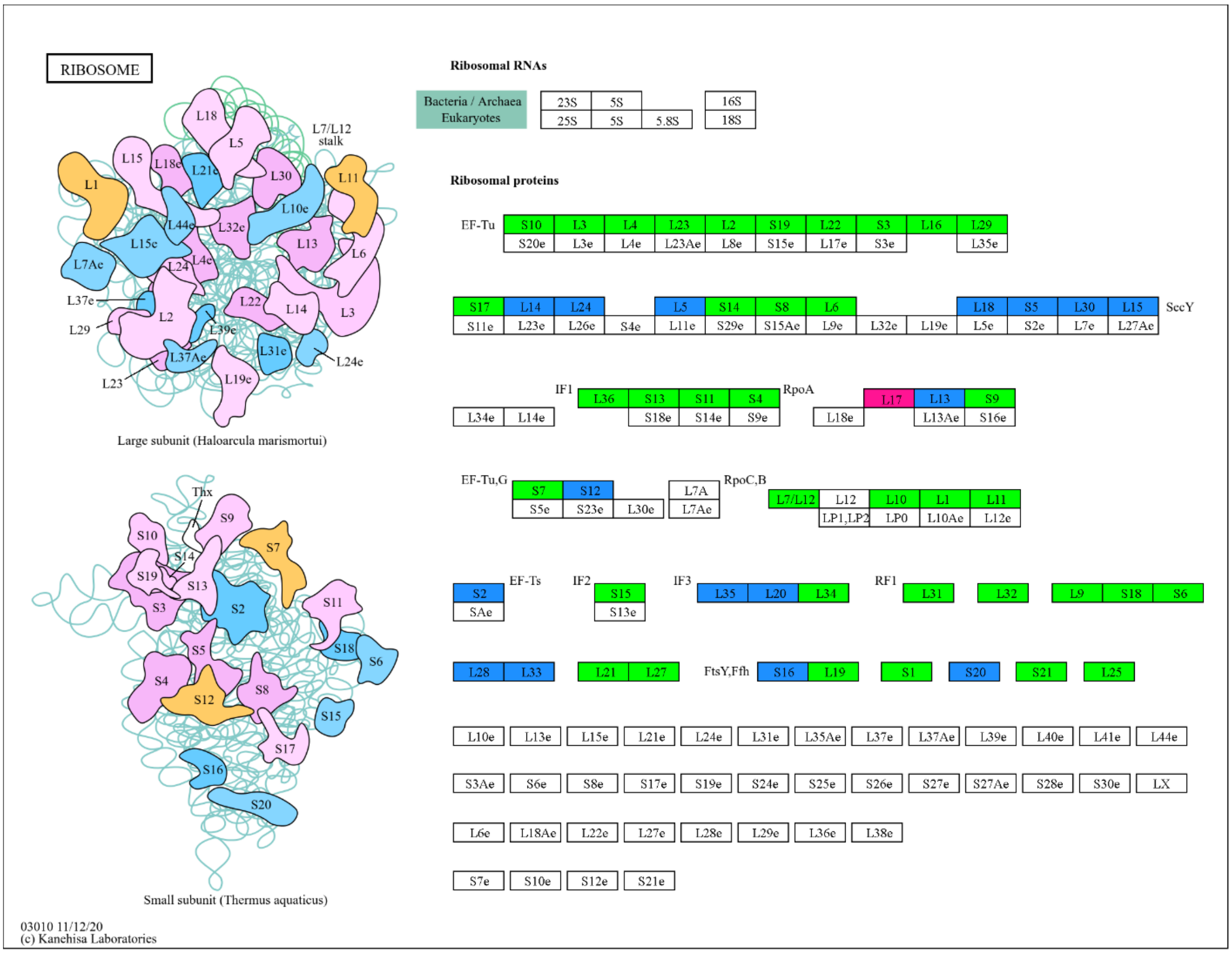


**Figure S23.** KEGG pathway for ribosome. Green represents enzymes/pathways not significantly enriched in iRNA and eRNA, light blue represents enzymes/pathways significantly enriched in iRNA while pink represents enzymes/pathways significantly enriched in eRNA (with p-adjusted < 0.01 and $|\log_2(\text{fold-change})| > 0.58$).

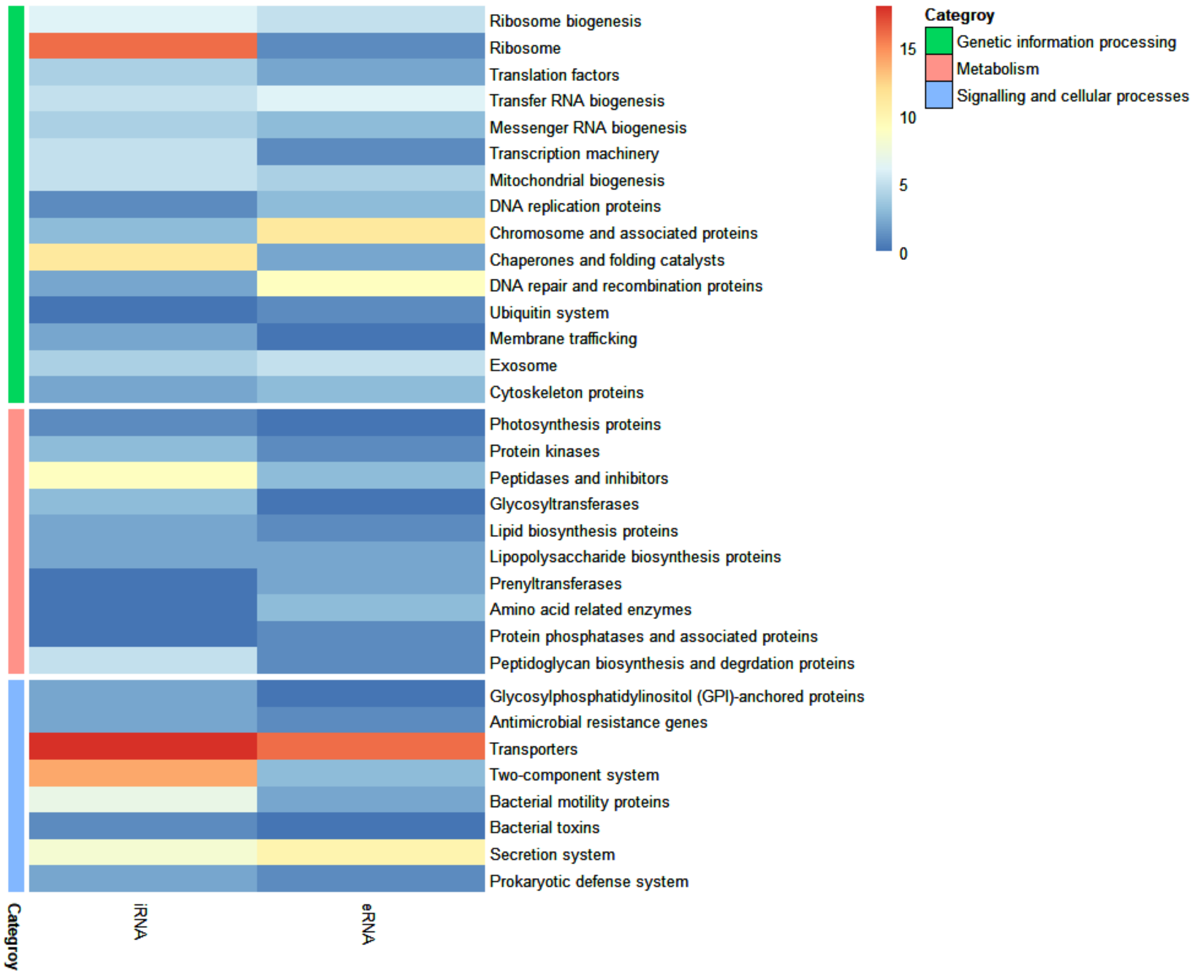


**Figure S24.** Heatmap showing the number of enriched genes for each BRITE category for eRNA and iRNA.

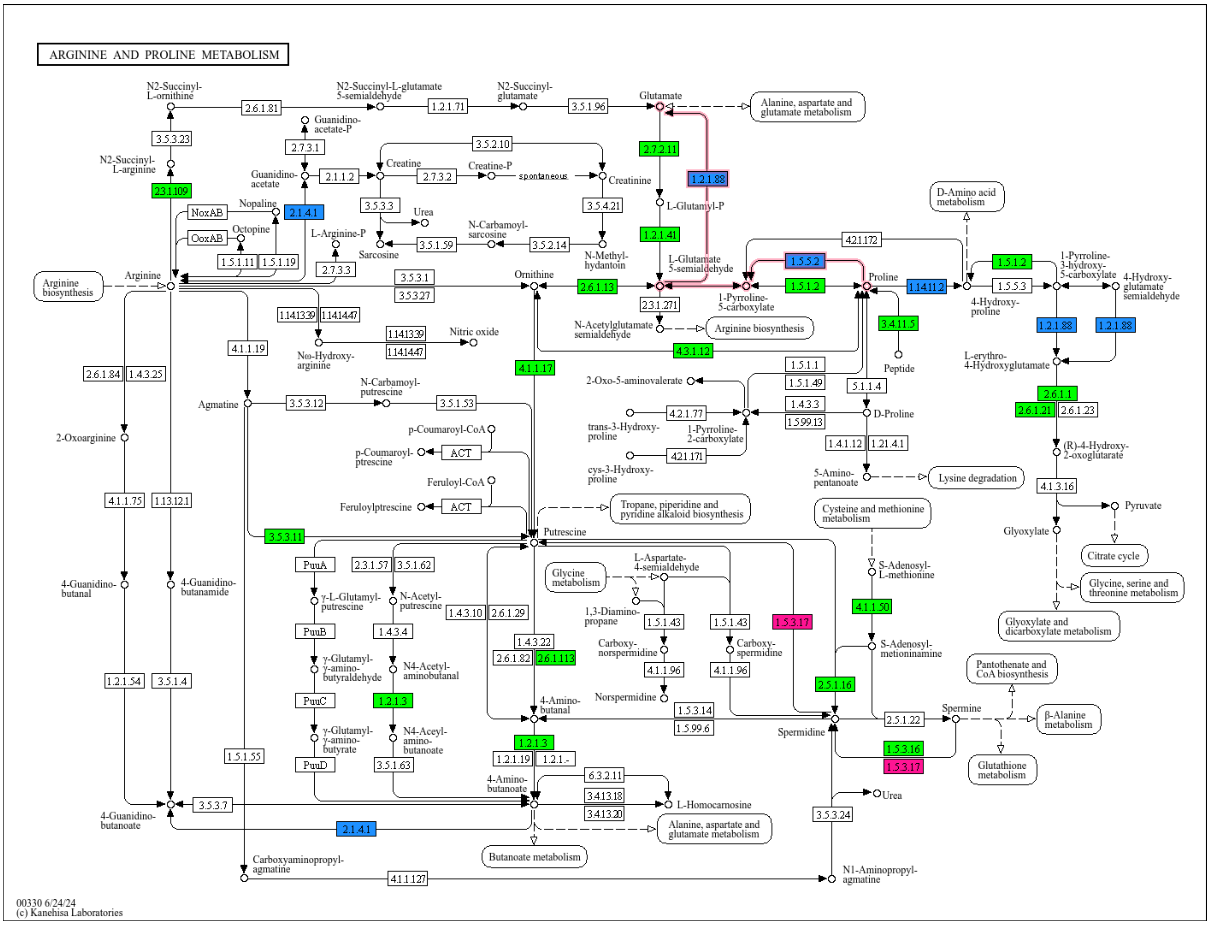


**Figure S25.** KEGG pathway for arginine and proline metabolism. Green represents enzymes/pathways not significantly enriched in iRNA and eRNA, light blue represents enzymes/pathways significantly enriched in iRNA while pink represents enzymes/pathways significantly enriched in eRNA (with p-adjusted < 0.01 and $\log_2$foldchange > ± 0.58).

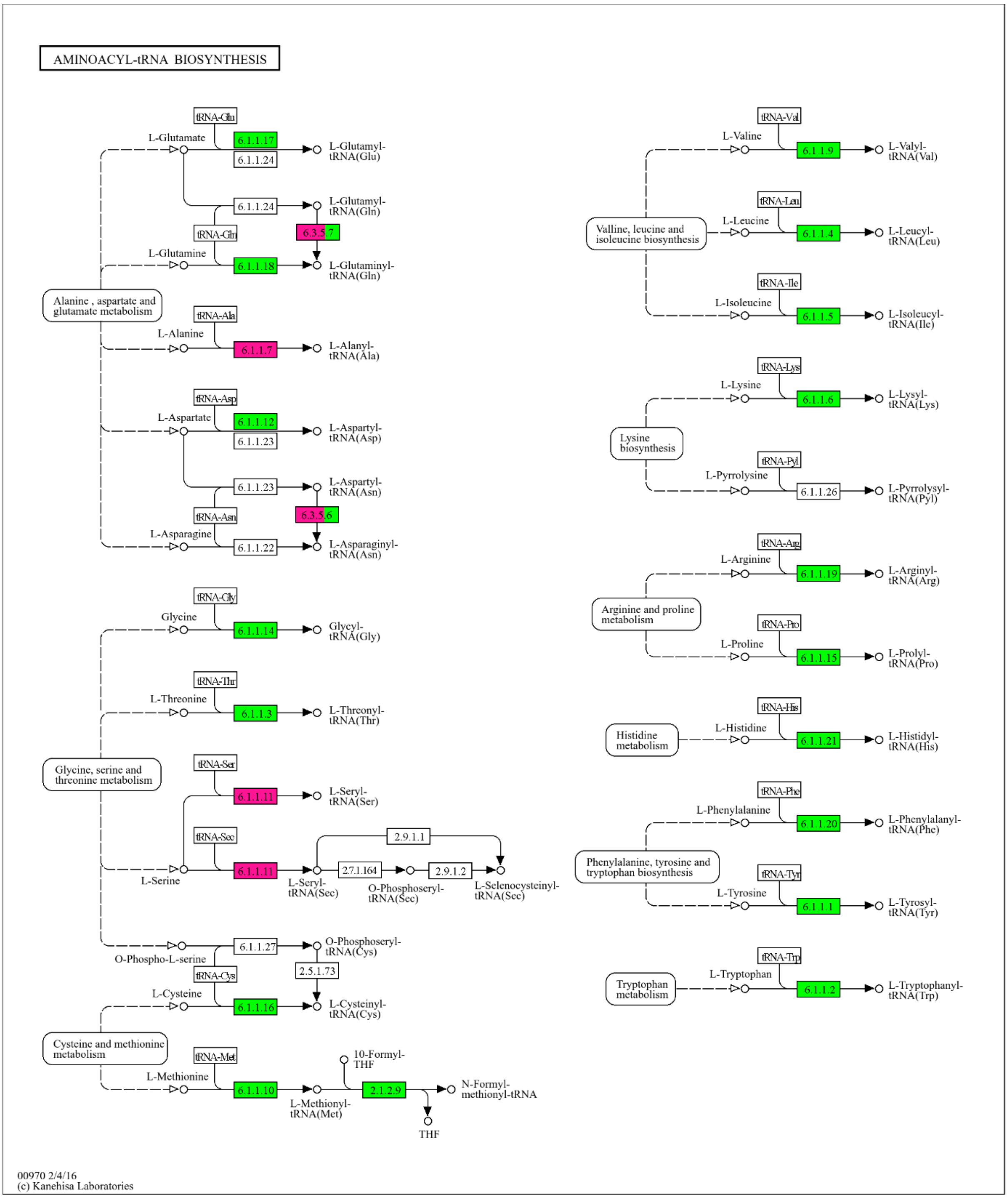


**Figure S26.** KEGG pathway for aminoacyl tRNA biosynthesis. Green represents enzymes/pathways not significantly enriched in iRNA and eRNA, light blue represents enzymes/pathways significantly enriched in iRNA while pink represents enzymes/pathways significantly enriched in eRNA (with p-adjusted < 0.01 and $\log_2$foldchange > ± 0.58).

**Table S1.** Finished water parameters at the tap measured at the end of the sampling process.

| Parameters | Quantity |
|---|---|
| Total organic carbon (mg/L) | 1.28 ± 0.03 |
| Total Carbon (mg/L) | 12.58 ± 0.06 |
| Inorganic carbon (mg/L) | 11.30 ± 0.04 |
| Total Nitrogen (mg/L) | 0.57 ± 0.08 |
| Monochloramine (mg/L $Cl_2$) | 0.90 ± 0.06 |
| Nitrite (mg/L N-$NO_2$) | 0.03 ± 0.001 |
| Nitrate (mg/L N-$NO_2$) | 0.25 ± 0.01 |
| Ammonia (mg/L N-$NH_3$) | 0.22 ± 0.001 |
| Chlorine (mg/L $Cl_2$) | 0.66 ± 0.001 |
| pH | 7.86 ± 0.04 |

**Table S2.** Concentration of purified nucleic acids submitted for sequencing. N.D indicates nucleic acid concentrations below detection limit.

| Replicate\ NAC Type | 1 | 2 | 3 | 4 |
|---|---|---|---|---|
| iDNA (ng/µl) | 17.5 | 18.2 | 13.7 | 21.8 |
| eDNA (ng/µl) | 2.4 | 1.9 | 3 | 2.2 |
| iRNA (ng/µl) | 55.1 | 98 | 77 | 13.3 |
| eRNA (ng/µl) | 57.2 | 48.8 | 55.9 | 38 |
| Total volume filtered (L) | 45.0 | 44.2 | 42.8 | 43.5 |
| Extraction blank | N.D | N.D | N.D | N.D |

**Table S3. Read classification summary by Kraken2 generated using Pavian. Total reads represent pair-end sequencing.**

| Name | Number of raw reads | Classified reads | Chordate reads | Artificial reads | Microbial reads | Bacterial reads | Viral reads | Fungal reads | Protozoan reads |
|---|---|---|---|---|---|---|---|---|---|
| eDNA1 | 1,754,599 | 54.40% | 0.14% | 0% | 54.30% | 54% | 0.00% | 0% | 0% |
| eDNA2 | 1,872,041 | 55.30% | 0.16% | 0% | 55.10% | 54.80% | 0.00% | 0% | 0% |
| eDNA3 | 1,699,794 | 54.90% | 0.16% | 0% | 54.80% | 54.50% | 0.00% | 0% | 0% |
| eDNA4 | 1,628,436 | 55% | 0.16% | 0% | 54.80% | 54.50% | 0.00% | 0% | 0% |
| eRNA1 | 36,942,284 | 98.70% | 0.02% | 0% | 98.70% | 98.60% | 0.00% | 0% | 0% |
| eRNA2 | 39,010,844 | 98.80% | 0.02% | 0% | 98.70% | 98.60% | 0.00% | 0% | 0% |
| eRNA3 | 35,477,155 | 98.80% | 0.02% | 0% | 98.70% | 98.60% | 0.00% | 0% | 0% |
| eRNA4 | 40,852,235 | 98.70% | 0.02% | 0% | 98.70% | 98.60% | 0.00% | 0% | 0% |
| iDNA1 | 1,663,903 | 55.50% | 0.10% | 0% | 55.40% | 55.20% | 0.00% | 0% | 0% |
| iDNA2 | 1,829,553 | 55% | 0.11% | 0% | 54.90% | 54.70% | 0.00% | 0% | 0% |
| iDNA3 | 1,703,798 | 55.60% | 0.11% | 0% | 55.50% | 55.30% | 0.01% | 0% | 0% |
| iDNA4 | 1,702,876 | 56% | 0.10% | 0% | 55.90% | 55.70% | 0.00% | 0% | 0% |
| iRNA1 | 32,959,886 | 98.70% | 0.01% | 0% | 98.70% | 98.60% | 0.00% | 0% | 0% |
| iRNA2 | 42,690,715 | 98.70% | 0.01% | 0% | 98.70% | 98.50% | 0.00% | 0% | 0% |
| iRNA3 | 36,333,977 | 98.60% | 0.01% | 0% | 98.60% | 98.50% | 0.00% | 0% | 0% |
| iRNA4 | 35,324,944 | 98.70% | 0.01% | 0% | 98.70% | 98.60% | 0.00% | 0% | 0% |

**Table S4.** β-diversity measures for community detected by intracellular and extracellular DNA and RNA based on the relative abundance of genera. $R^2$ and $p_{adonis}$ represent output for PERMANOVA implemented using 'adonis2()' command from the "vegan" package in R. $p_{permutest}$ represent significance in dispersion within each compared group.

| NAC | Distance | $R^2$ | $p_{adonis}$ | $p_{permutest}$ |
|---|---|---|---|---|
| **DNA** | Bray | 0.73 | 0.03* | 0.033* |
| **RNA** | Bray | 0.91 | 0.03* | 0.39 |

**Table S5. Top 30 COGs in DNA TPM values**

| COG | iDNA 1 | iDNA 2 | iDNA 3 | iDNA 4 | eDNA 1 | eDNA 2 | eDNA 3 | eDNA 4 | Annotation |
|---|---|---|---|---|---|---|---|---|---|
| ENOG410XNMH | 72275 | 79095 | 73894 | 74378 | 74659 | 79417 | 71720 | 68157 | Histidine kinase |
| ENOG4111JFI | 23366 | 25442 | 24169 | 25260 | 34010 | 36619 | 31107 | 29793 | Transposase |
| COG3696 | 19383 | 21410 | 20049 | 20144 | 19785 | 21029 | 18989 | 18303 | Putative silver efflux pump |
| COG3039 | 17940 | 19252 | 18395 | 17987 | 16823 | 17275 | 15646 | 15043 | Transposase and inactivated derivatives, IS5 family |
| COG1262 | 15398 | 16364 | 15278 | 15378 | 14004 | 14829 | 13212 | 12909 | Uncharacterized conserved protein |
| COG0610 | 12666 | 13543 | 12771 | 13022 | 12808 | 13764 | 11977 | 11658 | Type I site-specific restriction-modification system, R (restriction) subunit and related helicases |
| COG2931 | 10901 | 11648 | 11030 | 11234 | 11797 | 12460 | 11178 | 10702 | RTX toxins and related Ca2+-binding proteins |
| COG2826 | 9863 | 10722 | 10024 | 10241 | 12453 | 13194 | 11853 | 11758 | Transposase and inactivated derivatives, IS30 family |
| COG3436 | 9724 | 10018 | 9855 | 9709 | 12184 | 13320 | 12078 | 11734 | Transposase and inactivated derivatives |
| COG0840 | 10844 | 11988 | 11140 | 11072 | 10508 | 11126 | 10153 | 9722 | Methyl-accepting chemotaxis protein |
| COG3335 | 9243 | 10041 | 9533 | 9354 | 11928 | 12999 | 11294 | 11502 | Transposase and inactivated derivatives |
| COG1629 | 10460 | 11109 | 10429 | 10477 | 10308 | 11022 | 10198 | 9556 | Outer membrane receptor proteins, mostly Fe transport |
| COG0286 | 8956 | 9328 | 9026 | 9260 | 9180 | 9966 | 8859 | 8480 | Type I restriction-modification system methyltransferase subunit |
| COG0210 | 9026 | 9911 | 9006 | 9149 | 8852 | 9630 | 8610 | 8147 | Superfamily I DNA and RNA helicases |
| COG1132 | 8867 | 9682 | 8927 | 9081 | 9006 | 9510 | 8577 | 8194 | ABC-type multidrug transport system, ATPase and permease components |
| COG2801 | 8363 | 8846 | 8206 | 8520 | 8318 | 9009 | 7878 | 7705 | Transposase and inactivated derivatives |
| COG0500 | 7830 | 8558 | 8287 | 8320 | 7733 | 8338 | 7890 | 7296 | SAM-dependent methyltransferases |
| COG3505 | 7901 | 8746 | 8011 | 7908 | 7864 | 8483 | 7634 | 7298 | Type IV secretory pathway, VirD4 components |
| COG0488 | 7694 | 8138 | 7861 | 7780 | 7857 | 8537 | 7737 | 7628 | ATPase components of ABC transporters with duplicated ATPase domains |
| ENOG410XRFS | 7359 | 7769 | 7487 | 7474 | 8265 | 8426 | 7574 | 7526 | ATpase involved in D repair |
| COG0577 | 7524 | 8133 | 7624 | 7779 | 7487 | 8093 | 7128 | 7152 | ABC-type antimicrobial peptide transport system, permease component |
| COG0553 | 7010 | 7538 | 7036 | 7089 | 7445 | 8031 | 7357 | 7199 | Superfamily II DNA/RNA helicases, SNF2 family |
| COG3451 | 6915 | 7708 | 6987 | 6724 | 7362 | 7736 | 7052 | 6791 | Type IV secretory pathway, VirB4 components |
| COG0732 | 6586 | 7296 | 6974 | 6632 | 6898 | 7445 | 6616 | 6389 | Restriction endonuclease S subunits |

| | | | | | | | | | |
|---|---|---|---|---|---|---|---|---|---|
| COG3547 | 6162 | 6609 | 6314 | 6179 | 7482 | 8001 | 7081 | 6980 | Transposase and inactivated derivatives |
| ENOG410XR5J | 6369 | 7024 | 6607 | 6714 | 7172 | 7431 | 6829 | 6466 | Efflux transporter rnd family, mfp subunit |
| COG1639 | 6744 | 7192 | 6730 | 6880 | 6588 | 7204 | 6516 | 6166 | Predicted signal transduction protein |
| COG2197 | 5835 | 6764 | 6233 | 6057 | 7228 | 7756 | 6579 | 6474 | Response regulator containing a CheY-like receiver domain and an HTH DNA-binding domain |
| COG0784 | 6311 | 7091 | 6417 | 6588 | 6629 | 7150 | 6403 | 6030 | FOG: CheY-like receiver |
| COG4771 | 7041 | 7127 | 6757 | 6734 | 6269 | 6683 | 5982 | 5824 | Outer membrane receptor for ferrienterochelin and colicins |

**Table S6. KO in DNA. TPM values**

| KO | Function | iDNA_1 | iDNA_2 | iDNA_3 | iDNA_4 | eDNA_1 | eDNA_2 | eDNA_3 | eDNA_4 |
|---|---|---|---|---|---|---|---|---|---|
| K07488 | transposase | 23756 | 25900 | 24514 | 25666 | 35009 | 37661 | 32020 | 30607 |
| K01153 | type I restriction enzyme, R subunit [EC:3.1.21.3] | 16808 | 18424 | 17105 | 17554 | 17239 | 18701 | 16167 | 15987 |
| K07481 | transposase, IS5 family | 17874 | 19171 | 18346 | 17913 | 16841 | 17219 | 15590 | 14978 |
| K15726 | cobalt-zinc-cadmium resistance protein CzcA | 16126 | 18023 | 16766 | 16711 | 16809 | 17810 | 15972 | 15382 |
| K02014 | iron complex outermembrane recepter protein | 14116 | 14511 | 13740 | 13654 | 12517 | 13605 | 12595 | 11900 |
| K07484 | transposase | 11380 | 11938 | 11435 | 11301 | 13980 | 15120 | 13640 | 13333 |
| K03320 | ammonium transporter, Amt family | 12007 | 12835 | 12119 | 12317 | 11916 | 12978 | 11465 | 11045 |
| K21023 | diguanylate cyclase [EC:2.7.7.65] | 11760 | 12988 | 12313 | 12412 | 11330 | 12285 | 11079 | 10608 |
| K02004 | putative ABC transport system permease protein | 11005 | 12047 | 11200 | 11444 | 11003 | 11837 | 10490 | 10343 |
| K06919 | putative DNA primase/helicase | 10200 | 11476 | 10484 | 10479 | 10190 | 10729 | 9863 | 9355 |
| K03427 | type I restriction enzyme M protein [EC:2.1.1.72] | 9652 | 10117 | 9803 | 9962 | 10000 | 10907 | 9728 | 9269 |
| K13924 | two-component system, chemotaxis family, CheB/CheR fusion protein [EC:2.1.1.80 3.1.1.61] | 9557 | 10333 | 9845 | 9907 | 9744 | 10106 | 9654 | 8839 |
| K07482 | transposase, IS30 family | 8373 | 9056 | 8418 | 8648 | 10419 | 11100 | 9986 | 9852 |
| K07497 | putative transposase | 9356 | 10021 | 9364 | 9605 | 9415 | 9975 | 8829 | 8573 |
| K15727 | membrane fusion protein, cobalt-zinc-cadmium efflux system | 7725 | 8502 | 7946 | 7977 | 8445 | 8862 | 8091 | 7702 |
| K03657 | DNA helicase II / ATP-dependent DNA helicase PcrA [EC:3.6.4.12] | 7839 | 8707 | 7821 | 8038 | 7871 | 8544 | 7661 | 7285 |
| K03655 | ATP-dependent DNA helicase RecG [EC:3.6.4.12] | 7681 | 8468 | 7725 | 7765 | 7395 | 8051 | 7090 | 7040 |
| K11912 | serine/threonine-protein kinase PpkA [EC:2.7.11.1] | 7869 | 8172 | 7990 | 8162 | 7425 | 7722 | 6964 | 6707 |
| K01338 | ATP-dependent Lon protease [EC:3.4.21.53] | 7109 | 7733 | 7407 | 7397 | 7725 | 8532 | 7339 | 7267 |
| K03205 | type IV secretion system protein VirD4 | 7421 | 8263 | 7471 | 7443 | 7314 | 7764 | 7018 | 6696 |
| K01156 | type III restriction enzyme [EC:3.1.21.5] | 7067 | 7620 | 7087 | 7332 | 7253 | 7575 | 7134 | 6654 |
| K20530 | type IV secretion system protein TrbE | 6915 | 7684 | 6995 | 6715 | 7415 | 7786 | 7083 | 6808 |
| K06147 | ATP-binding cassette, subfamily B, bacterial | 6921 | 7683 | 7102 | 7349 | 6785 | 7307 | 6617 | 6247 |
| K07486 | transposase | 6120 | 6602 | 6288 | 6171 | 7458 | 7970 | 7029 | 6940 |

| | | | | | | | | | |
|---|---|---|---|---|---|---|---|---|---|
| K01077 | alkaline phosphatase [EC:3.1.3.1] | 6777 | 7362 | 7101 | 6793 | 6631 | 6878 | 6400 | 5916 |
| K03406 | methyl-accepting chemotaxis protein | 6721 | 7129 | 6724 | 6661 | 6191 | 6685 | 6176 | 5920 |
| K07093 | uncharacterized protein | 6611 | 6592 | 6599 | 6412 | 6410 | 6767 | 6240 | 5782 |
| K03043 | DNA-directed RNA polymerase subunit beta [EC:2.7.7.6] | 6178 | 7047 | 6618 | 6312 | 5466 | 5805 | 5532 | 5021 |
| K01768 | adenylate cyclase [EC:4.6.1.1] | 5687 | 6291 | 6010 | 6055 | 5950 | 6534 | 5742 | 5567 |
| K01126 | glycerophosphoryl diester phosphodiesterase [EC:3.1.4.46] | 5894 | 6418 | 6147 | 6138 | 5771 | 6181 | 5793 | 5408 |

**Table S7. COG in RNA. TPM values**

| COG | iRNA 1 | iRNA 2 | iRNA 3 | iRNA 4 | eRNA 1 | eRNA 2 | eRNA 3 | eRNA 4 | Annotation |
|---|---|---|---|---|---|---|---|---|---|
| ENOG41125ZN | 267224 | 358159 | 331090 | 214812 | 261154 | 242740 | 219872 | 267405 | Unknown Function |
| COG2197 | 179414 | 248953 | 227849 | 159940 | 199504 | 201546 | 172664 | 214078 | Response regulator containing a CheY-like receiver domain and an HTH DNA-binding domain |
| COG4775 | 174313 | 241679 | 222559 | 126164 | 157568 | 146977 | 123323 | 153714 | Outer membrane protein/protective antigen OMA87 |
| ENOG410XPHF | 140949 | 175903 | 162237 | 136726 | 136031 | 127686 | 127093 | 149532 | Efflux transporter rnd family, mfp subunit |
| ENOG410Y13W | 92294 | 105311 | 92733 | 103024 | 133363 | 116197 | 119718 | 124233 | PG1 protein, homology to Homo sapiens |
| COG1053 | 97366 | 124718 | 117633 | 105639 | 84868 | 92034 | 83255 | 100145 | Succinate dehydrogenase/fumarate reductase, flavoprotein subunit |
| COG5563 | 77334 | 105710 | 94798 | 68383 | 71926 | 83470 | 69488 | 90293 | Predicted integral membrane proteins containing uncharacterized repeats |
| ENOG410XSVV | 83231 | 105885 | 91445 | 95463 | 49616 | 51355 | 51392 | 61423 | Methane monooxygenase ammonia monooxygenase, subunit C |
| ENOG410Y1V8 | 79853 | 101180 | 87516 | 59073 | 62859 | 58691 | 48186 | 65417 | Unknown Function |
| ENOG410Y0ZG | 59201 | 73424 | 61383 | 75476 | 49325 | 49640 | 53126 | 61531 | monooxygenase, subunit B |
| ENOG4111MT1 | 54289 | 74814 | 63659 | 66553 | 54802 | 53489 | 50009 | 64996 | cytochrome p460 |
| ENOG4110CTU | 63771 | 84870 | 71865 | 72171 | 39644 | 42018 | 40921 | 51057 | methane monooxygenase ammonia monooxygenase subunit A |
| ENOG410XS4Q | 42037 | 54587 | 46996 | 32841 | 46260 | 40062 | 37685 | 44708 | Short-chain dehydrogenase reductase Sdr |
| COG1792 | 26311 | 33100 | 32193 | 22210 | 33935 | 29280 | 27383 | 30674 | Cell shape-determining protein |
| COG2132 | 19736 | 26145 | 22833 | 26229 | 18861 | 18358 | 18908 | 22285 | Putative multicopper oxidases |
| ENOG4111IGE | 17633 | 20693 | 19268 | 18778 | 20493 | 19299 | 20922 | 25352 | Unknown Function |
| COG0464 | 19283 | 26281 | 24564 | 14039 | 17329 | 17504 | 14129 | 18193 | ATPases of the AAA+ class |
| ENOG410XQ4U | 14730 | 17834 | 16562 | 13559 | 18288 | 15760 | 16602 | 17653 | (LipO)protein |
| COG0607 | 12271 | 13480 | 13158 | 17959 | 15139 | 16188 | 20792 | 20244 | Rhodanese-related sulfurtransferase |
| ENOG410XPVG | 15056 | 19158 | 16145 | 18214 | 13346 | 14157 | 14109 | 16582 | hydroxylamine oxidase |
| COG1262 | 15154 | 21134 | 19478 | 13018 | 10415 | 10294 | 10161 | 10755 | Uncharacterized conserved protein |

| | | | | | | | | | |
|---|---|---|---|---|---|---|---|---|---|
| ENOG4111UQ9 | 12809 | 18070 | 15821 | 14569 | 11409 | 11240 | 10170 | 12760 | cytochrome C, class I |
| COG0628 | 8411 | 8912 | 9154 | 13262 | 13074 | 14585 | 16749 | 18616 | Predicted permease |
| COG0058 | 12639 | 15560 | 14445 | 11655 | 12543 | 11250 | 11319 | 12679 | Glucan phosphorylase |
| COG2931 | 12860 | 16092 | 13903 | 14833 | 7149 | 7347 | 7161 | 8730 | RTX toxins and related Ca2+-binding proteins |
| COG1622 | 12494 | 15849 | 13599 | 14219 | 6306 | 6516 | 6480 | 7983 | Heme/copper-type cytochrome/quinol oxidases, subunit 2 |
| COG0843 | 10323 | 13015 | 10940 | 13550 | 7787 | 8027 | 8249 | 10109 | Heme/copper-type cytochrome/quinol oxidases, subunit 1 |
| ENOG410Z8CT | 9400 | 11247 | 9678 | 12951 | 6319 | 6641 | 6760 | 7917 | PEP-CTERM motif |
| ENOG410Y4NW | 9061 | 11318 | 9942 | 10319 | 3618 | 3623 | 3584 | 4264 | Porin, Gram-negative type |
| ENOG410XURY | 6262 | 8154 | 7137 | 7589 | 5889 | 6011 | 5804 | 6945 | Unknown Function |
| ENOG41125ZN | 267224 | 358159 | 331090 | 214812 | 261154 | 242740 | 219872 | 267405 | Unknown Function |
| COG2197 | 179414 | 248953 | 227849 | 159940 | 199504 | 201546 | 172664 | 214078 | Response regulator containing a CheY-like receiver domain and an HTH DNA-binding domain |

### Table S8. Top 30 KOs in RNA TPM values

| KO | Function | iRNA_1 | iRNA_2 | iRNA_3 | iRNA_4 | eRNA_1 | eRNA_2 | eRNA_3 | eRNA_4 |
|---|---|---|---|---|---|---|---|---|---|
| K07277 | outer membrane protein insertion porin family | 174317 | 241681 | 222559 | 126168 | 157572 | 146977 | 123323 | 153716 |
| K15727 | membrane fusion protein, cobalt-zinc-cadmium efflux system | 141042 | 175991 | 162377 | 136991 | 136191 | 127812 | 127259 | 149793 |
| K00239 | succinate dehydrogenase / fumarate reductase, flavoprotein subunit [EC:1.3.5.1 1.3.5.4] | 97366 | 124718 | 117633 | 105639 | 84868 | 92034 | 83255 | 100143 |
| K10946 | methane/ammonia monooxygenase subunit C | 83428 | 106067 | 91642 | 95638 | 49687 | 51452 | 51472 | 61495 |
| K10945 | methane/ammonia monooxygenase subunit B | 59634 | 73802 | 61817 | 75972 | 49830 | 50093 | 53584 | 62118 |
| K10944 | methane/ammonia monooxygenase subunit A [EC:1.14.18.3 1.14.99.39] | 63778 | 84887 | 71875 | 72186 | 39650 | 42028 | 40930 | 51065 |
| K00059 | 3-oxoacyl-[acyl-carrier protein] reductase [EC:1.1.1.100] | 42136 | 54749 | 47105 | 33018 | 46461 | 40224 | 37878 | 44923 |
| K08738 | cytochrome c | 27963 | 38711 | 32826 | 34747 | 26268 | 26464 | 24998 | 31773 |
| K03570 | rod shape-determining protein MreC | 26311 | 33100 | 32193 | 22210 | 33935 | 29280 | 27383 | 30674 |
| K00368 | nitrite reductase (NO-forming) [EC:1.7.2.1] | 19115 | 25412 | 22240 | 25557 | 18240 | 17775 | 18320 | 21619 |
| K10535 | hydroxylamine dehydrogenase [EC:1.7.2.6] | 15056 | 19158 | 16145 | 18214 | 13346 | 14157 | 14109 | 16582 |
| K00688 | glycogen phosphorylase [EC:2.4.1.1] | 12639 | 15560 | 14445 | 11655 | 12543 | 11250 | 11319 | 12679 |
| K02275 | cytochrome c oxidase subunit II [EC:1.9.3.1] | 12446 | 15817 | 13557 | 14187 | 6245 | 6458 | 6446 | 7921 |
| K02274 | cytochrome c oxidase subunit I [EC:1.9.3.1] | 10322 | 13005 | 10933 | 13548 | 7776 | 8021 | 8247 | 10105 |
| K20533 | type IV secretion system protein TrbI | 6636 | 8141 | 8128 | 6253 | 5720 | 6025 | 4988 | 6438 |
| K02276 | cytochrome c oxidase subunit III [EC:1.9.3.1] | 5753 | 7446 | 6657 | 6659 | 4492 | 4927 | 4457 | 5814 |
| K03798 | cell division protease FtsH [EC:3.4.24.-] | 5151 | 6528 | 5773 | 6097 | 3537 | 3439 | 3484 | 4183 |
| K02569 | cytochrome c-type protein NapC | 3780 | 5605 | 4690 | 3957 | 4568 | 4808 | 4307 | 5306 |
| K02358 | elongation factor Tu | 4702 | 6666 | 5554 | 4909 | 3235 | 3407 | 3343 | 3879 |
| K02258 | cytochrome c oxidase assembly protein subunit 11 | 4330 | 5769 | 4994 | 4727 | 3290 | 3325 | 3141 | 3898 |
| K01077 | alkaline phosphatase [EC:3.1.3.1] | 4384 | 5223 | 4880 | 5883 | 2918 | 2955 | 2870 | 3499 |
| K07755 | arsenite methyltransferase [EC:2.1.1.137] | 4354 | 5305 | 5549 | 3676 | 3366 | 3311 | 3023 | 3791 |

| | | | | | | | | | |
|---|---|---|---|---|---|---|---|---|---|
| K01601 | ribulose-bisphosphate carboxylase large chain [EC:4.1.1.39] | 3290 | 4098 | 3305 | 3771 | 3161 | 3259 | 3168 | 3847 |
| K02112 | F-type H+/Na+-transporting ATPase subunit beta [EC:7.1.2.2 7.2.2.1] | 3560 | 4807 | 4245 | 4036 | 2501 | 2862 | 2658 | 3180 |
| K01607 | 4-carboxymuconolactone decarboxylase [EC:4.1.1.44] | 3015 | 4040 | 3502 | 3803 | 2855 | 2774 | 2688 | 3262 |
| K03046 | DNA-directed RNA polymerase subunit beta' [EC:2.7.7.6] | 2115 | 2843 | 2486 | 2787 | 2598 | 2557 | 2425 | 2983 |
| K06137 | pyrroloquinoline-quinone synthase [EC:1.3.3.11] | 2025 | 2579 | 2177 | 2655 | 2632 | 2549 | 2557 | 3324 |
| K03088 | RNA polymerase sigma-70 factor, ECF subfamily | 2775 | 3905 | 3248 | 2813 | 1705 | 1753 | 1360 | 1887 |
| K00549 | 5-methyltetrahydropteroyltriglutamate--homocysteine methyltransferase [EC:2.1.1.14] | 1628 | 1969 | 1724 | 1898 | 2553 | 2575 | 2489 | 3069 |
| K15726 | cobalt-zinc-cadmium resistance protein CzcA | 1614 | 2139 | 1908 | 2245 | 2146 | 2183 | 1975 | 2496 |
